# Long-Term Organ Culture Reveals Differential Stem Cell–Driven Remodeling in Myometrium and *MED12*-Mutant Uterine Leiomyoma

**DOI:** 10.1101/2025.08.19.670360

**Authors:** Paula Vázquez, Ana Salas, Silvia Beltrán-Flores, Francisco Monte de Oca, Araceli Delgado, Teresa A. Almeida

## Abstract

Uterine leiomyomas or fibroids are highly prevalent benign tumors of the female reproductive tract, often causing significant symptoms and requiring surgical intervention, leading to substantial healthcare costs worldwide. Their molecular pathogenesis remains incompletely understood, but evidence suggests that somatic stem cells play a pivotal role in myometrial growth, whereas a genetic alteration, particularly mutations in *MED12,* may transform a myometrial stem cell into a tumor-initiating cell, promoting fibroid growth.

In organ cultures of fibroids and myometrium, most differentiated cells degenerated by day 7, whereas stem cells remained quiescent and viable within their native niches. Notably, between days 15 and 29, hypoxia-induced activation triggered stem cell proliferation and differentiation within the *ex vivo* slices.

Transcriptomic profiling revealed statistically significant upregulation of stemness-associated genes, including *HMGA*, *ITG*, *KLF*, *HOX*, and *SOX* family members, in long-term cultured slices compared with baseline tissue, and between normal and tumor cultures. Reactome pathway enrichment analysis further identified distinct metabolic, extracellular matrix remodeling, immune surveillance, angiogenic, and cell death– related programs distinguishing myometrial from leiomyoma cultures. Furthermore, previously reported gene and pathway differences between healthy and fibroid tissues were robustly confirmed, validating the culture model.

In conclusion, our findings establish long-term organ culture as a powerful, physiologically relevant platform for investigating stem cell dynamics in myometrium and uterine leiomyoma. They also provide proof of concept for extending this approach to other tissue types, enabling the discovery of mechanisms underlying stem cell activation, differentiation, and death, with broad translational potential in regenerative medicine and cancer biology.

## INTRODUCTION

Uterine leiomyomas (UL), also called myomas or fibroids, are benign tumors of the female genital tract, affecting approximately 70% of Caucasian and 80% of Black African women by age 50^1^. Roughly 30–50% of fibroids cause symptoms that significantly impact quality of life, with heavy or prolonged menstrual bleeding being the most common and the primary indication for surgery. Treatment typically involves myomectomy for women wishing to preserve fertility, or hysterectomy for those who have completed childbearing. Although minimally invasive alternatives exist, they are not widely adopted, and surgery remains the standard approach, contributing to substantial healthcare costs worldwide^1–3^.

Although the etiology of uterine leiomyomas remains unclear, transcriptomic analyses have identified distinct molecular subgroups: UL harboring mutations in mediator complex subunit 12, *MED12*, rearrangements in high mobility group AT-hook 2, *HMGA2*, biallelic inactivation of fumarate hydratase, *FH*, deletions of collagen, type IV, alpha 5, and collagen, type IV, alpha 6, *COL4A5–COL4A6*, and those lacking all four driver alterations, termed quadruple-negative leiomyomas^4,5^. Mutations in *MED12* occur in around 70% of UL, constituting the most frequent subgroup, and in 99% of these tumors, the mutation occurs in exon 2^6^.

Somatic stem cells (SC) are essential in the female reproductive tract, where they promote remarkable plasticity and exceptional regenerative potential for uterine pregnancy-induced expansion^7,8^. A range of surface markers have been characterized in human myometrium (MM) stem/progenitor cells, including CRIP1, Stro-1/CD44, CD34 and CD49f (encoding by *ITGA6*)^9–11^. In UL, the most recent hypothesis regarding origin suggests that a genetic alteration, such as the *MED12* mutation, may transform a myometrial stem cell into a fibroid stem cell, initiating uncontrolled proliferation until the cell ultimately differentiates into a mature fibroid smooth muscle cell (SMC)^12,13^.

Supporting this concept, specific cell populations isolated from UL tissue that express the surface marker CD49b (encoding by *ITGA2*) and exhibit high levels of the stem cell transcription factors *OCT4*, *KLF4*, and *NANOG*, have demonstrated the capacity to form colonies *in vitro* and regenerate tumors *in vivo*, strongly indicating that CD49b⁺ cells possess stem cell–like properties^14^. In addition, UL smooth muscle cells exhibit higher expression of the CD24 surface marker (CD24^hi^) compared to normal myometrium^15^. These CD24^hi^ cells show characteristics of progenitor cells, including diminished alpha-smooth muscle actin expression and elevated levels of the stem cell surface marker CD73 (encoding by NT5E).

Increasing evidence points to low-oxygen levels as a condition that may stimulate tissue-specific stem cell survival and growth^16,17^. Notably, while myometrial SCs demonstrated no *in vitro* proliferation in a normoxic (20% O2) environment, they exhibited robust growth under 2% oxygen tension, successfully differentiating into SMCs. Recreating hypoxia in SC cultures is challenging and requires specific methods and specialized equipment. Additionally, cells grown on culture plates lack extracellular matrix (ECM) and neighbouring cells of the tissue microenvironment. We have previously set up an organ culture of UL and MM, where cells in tissue slices maintained their structural integrity and functional activity for seven days in culture.

Subsequent cell death led to a dramatic reduction in cells within the tissue sections. Surprisingly, after 15-20 days of culture, new SMCs repopulated almost the entire tissue slice^18^, which remained covered with cells until 29-30 days, after which cells disappear. Our study investigated transcriptomic changes in paired myometrial and fibroid tissues under long-term organ culture (LT-culture) conditions. Comparing each tissue at baseline, after surgery removal, with its LT-culture counterpart revealed gene expression programs associated with stem cell activation, proliferation, differentiation, and apoptosis. Additional tissue-specific differences emerged when myometrium and fibroid tissues were compared under LT-culture. As a proof of concept, we identified several genes and pathways previously reported to be dysregulated in fibroids, as well as novel genes with key roles in stem cell function.

## MATERIALS AND METHODS

### Collection of tissue samples

This study was approved by the Institutional Review Board of the Committee for Drug Research Ethics of the Complejo Hospitalario Universitario de Canarias (CHUC_2022_90, BioNanoGene2022). Informed consent was obtained from the patients before FMO collected any samples. All experiments involving human tissues were performed by the principles outlined in the Declaration of Helsinki.

Four Caucasian female patients aged 44–51 years, admitted to Hospital Quirónsalud Tenerife, were enrolled in this study between January and February 2023. The patients underwent hysterectomy for polymyomatous uterus, irregular and heavy menstrual bleeding, and bulk symptoms. Tumors analyzed were up to 5–8 cm in size, and, for each sample, their paired myometrium was also collected. The histopathological analysis using standard H&E staining performed by a pathologist indicated tumors with benign histology with no sign of malignancy, nuclear atypia, mitotic figures or necrosis. After surgery, samples were immediately submerged in sterile Hank’s balanced salt solution (HBSS) supplemented with 0.25 µg/mL amphotericin B, 100 U/ mL penicillin and 100 µg/mL streptomycin (Sigma-Aldrich Co., USA). Tissue pieces were then transported to the laboratory and handled under sterile conditions. The samples were processed within 2 hours after surgery.

### Tissue sectioning

UL and MM pieces were removed from the tubes and placed in Petri dishes containing sterile, cold HBSS supplemented with antibiotics and antifungal agents. Then, a section approximately 1 cm thick and 2-3 cm² was dissected and gently held between the forceps. Next, the tumor piece was cut in the middle using a double-edge carbon steel blade (Ted Pella Inc., USA) to obtain two thinner sections. The procedure was repeated with each section until the slice reached a thickness that allowed the serrated forceps tip to be seen through the tissue. Then, the slice was placed on the top of a glass plate with a 5 x 5 mm square mark to trim and obtain pieces of similar size. The tumor slices were kept well-soaked in cold HBSS during the procedure until they were cultured.

### Organ tissue culture

UL and MM tissue slices were cultivated onto polystyrene CELLSCAFLD® 3D (JET BIOFIL, China) and placed in a six-well culture plate. Four tissue slices were placed onto each scaffold, and 1 mL of Dulbecco’s modified Eagle’s medium (DMEM, Biowest, France) supplemented with 10% fetal bovine serum (FBS, Lonza, Spain), 100 U/mL penicillin, 100 µg/mL streptomycin and 2 mM L-Glutamine (Sigma-Aldrich, USA) was added to each well. A medium drop was added to the tissue slice to maintain the explant’s humidity. Tissue culture plates were maintained at 37 °C in a 5% CO2 humidified incubator on an orbital shaker (60 rpm). The medium was changed every day.

### Tissue histology characterization

To assess the morphological integrity of the tissue slices, two replicates of each tumor at days 0, 7, 15, 20, 25 and 29 were hematoxylin-eosin (H&E) stained. The tissue slices were painted with green Ink for biopsies (Green Ink, VWR Q-Path Chemicals, USA) in the air-contact side before their fixation to evaluate changes between both sides of the slice. Tissues were fixed in 10% buffered formalin, then embedded in paraffin in horizontal and vertical orientations to allow for different sectioning planes, and cut in 4-μm-thick sections. The sections were deparaffinized, hydrated, and H&E stained as described before^18^.

### RNA isolation

Tissue samples were placed into lysing matrix tubes D containing 500 µl of Tri-reagent (Zymo Research) and homogenized using FastPrep-24TM 5 G instrument (MP Biomedicals, Illkirch, France) twice for 30 s at a speed of 6 m/s. The sample tubes were placed on ice for 5 min between pulses. Then, chloroform was added to the lysate, centrifuged, and the RNA from the top aqueous phase was isolated and purified using Direct-zol RNA Microprep kit according to the manufacturer’s protocol (Zymo Research). Residual genomic DNA was removed by incubating the RNA samples with RNase-free DNase I and RNasin according to the manufacturer’s instructions (Promega Corp., Madison, WI, USA). The effectiveness of the DNase treatment was assessed in samples with no reverse transcriptase added (RT-negative). RNA was quantified by absorbance using a NanoDrop ND-1000 spectrophotometer (ThermoFisher Scientific, Waltham, MA, USA). Total RNA quality was assessed using an Agilent Bioanalyzer 2100. RNA integrity number (RIN) ranged from 3 to 7.5.

### MED12 mutation detection

Two tissue slices were analysed to confirm driver genetic alterations during the culture period, one at collection time (T0) and the other after LT-culture. Amplification of *MED12* cDNA was performed with primers located in exon 1 and exon 2, covering the hot spot region where 99% of mutations have been described. PCR products were cleaned up by ExoSAP treatment using Illustra™ ExoProStar™ 1-Step (GE Healthcare Biosciences, USA) following the manufacturer’s instructions. Sequencing reactions were performed for both strands at the Genomic Service of the University of La Laguna (SEGAI). For each sample, forward and reverse electropherograms were checked manually using Chromas 2.6.6 (Technelysium Pty Ltd., South Brisbane, Queensland, Australia 2018).

### HMGA2 expression

To detect fibroids with *HMGA2* overexpression, we performed PCR with primers previously designed^18^. PCR mixes contained 0.25 pmol of each primer, 1 unit of TEMPase Hot Start DNA Polymerase (VWR), 150 µM dNTPs, and 4 µL of cDNA (1/12 dilution) in a final volume of 20 µL. The cycling conditions were 95 °C for 15 min, followed by 40 cycles of 95 °C for 15 s, 66 °C for 20 s, and 72 °C for 30 s. For each experiment, a non-sample reaction and a positive control were included. The PCR products were separated by agarose gel electrophoresis, and the amplicon size was verified by comparison with a 100-bp DNA ladder.

### 3’ mRNA-Seq library preparation and sequencing

Given RNA quality obtained from LT-cultures (RIN 3–7.5), we selected a 3’-end RNA sequencing approach, which is more robust to RNA fragmentation and optimized for reliable gene expression quantification under such conditions^19^.

RNA-Seq library preparation and sequencing were outsourced to a commercial service provider (Seqplexing Multiplex SL, Valencia, Spain). The differential expression profiling study was conducted on 4 tumors and 4 matched myometria obtained after surgery (T0) and long-term culture (LT-culture), when cells repopulated the tissue slices (T-15, T-20, T-25 and T-29). The process begins with reverse transcription of 500 ng of total RNA using an oligo-dT primer that specifically binds to the poly(A) tail of mRNA, ensuring the selective capture of these transcripts. The RNA strand is then degraded, leaving a single-stranded cDNA. Next, second-strand synthesis is performed using random priming, generating a double-stranded cDNA library with incorporated unique molecular identifiers (UMIs) for further processing. Subsequently, a first PCR is carried out to amplify the cDNA library to increase the amount of genetic material available for analysis. Then, a second PCR is performed, in which an adapter is introduced to ensure that the cDNA fragments are properly anchored to the platform, allowing for an accurate reading of the sequences.

QIAxcel Advanced System assessed library quality to detect degradation or low concentration before sequencing. Sequencing was performed on the Illumina NovaSeq X with paired-end 2 × 150 bp reads. Raw FASTQ data were used for bioinformatic analysis.

### Data Analysis

The bioinformatics pipeline began with quality control and trimming of raw sequencing reads, removing adapters and poly(A) sequences. FastQC evaluated the quality of FASTQ files, ensuring good %Q20 scores and absence of errors. UMIs were processed using UMI-tools, relocating molecular markers to the read header for accurate mapping. Reads were aligned to the “GRCh38” reference genome with STAR, accounting for splicing junctions (v2.7.10)^20^. Duplicate reads were removed using UMIs to distinguish unique molecules. Gene expression was quantified using HTSeq-count and normalized to adjust for sequencing depth. Differential expression analysis was performed with DESeq2, identifying significant gene differences. PCA clustering in R evaluated sample quality, while visualization tools like ggplot and pheatmap facilitated data interpretation.

The expression of each gene was reported as the base 2 logarithm of the ratio of the value obtained when comparing the LT-culture of MM and UL to their corresponding T0 and the LT-culture of UL vs. the LT-culture of the matching MM. The cutoff values for log2 fold change (log2FC) were set at 1 for upregulated and −1 for downregulated genes. The Benjamini-Hochberg correction method for false discovery rate (FDR) was used to adjust resulting p-values for multiple testing with a cutoff of < 5% (q-value).

### Gene Ontology (GO) and KEGG pathway enrichment analysis

Gene Ontology (GO) and Kyoto Encyclopedia of Genes and Genomes (KEGG) pathway enrichment analyses were conducted to identify significantly affected biological functions and pathways associated with differentially expressed genes. Both analyses used adjusted p-values (p-adj) to assess statistical significance and calculated a Normalized Enrichment Score (NES) to quantify the direction (activation or repression) and magnitude of enrichment. GO analysis focused on categorizing changes within biological processes (BP), cellular components (CC), and molecular functions (MF), while KEGG analysis identified impacted metabolic and signaling pathways. Enriched terms and pathways with the most significant p-adj values were ranked and summarized based on the number of involved genes and their expression patterns, highlighting the most affected biological categories in the dataset.

### Reactome enrichment analysis

Pathway enrichment analysis was performed using the ReactomePA (v1.50.0)^21^ and clusterProfiler (v4.14.4)^22^ packages in R. The enrichPathway function was applied to identify significantly enriched pathways from the Reactome database, using all genes from the previous differential expression analysis as input. The results were visualized using the compareCluster function for grouped comparisons, and enrichment outcomes were displayed using the dotplot function.

## RESULTS

### Normal and *MED12*-mutant organ cultures repopulate with smooth muscle cells

After seven days in culture, we observed a progressive decrease in cells in MM and UL organ culture, until cells were almost completely absent. Surprisingly, after 15-20 days in culture, the tissue slices from the four fibroids and paired myometria were repopulated, with cells distributed throughout their surface. Immunostaining with desmin and vimentin demonstrated that most of them were SMC (Supplementary Figure S1).

We did not detect *HMAG2* expression in any of the tumors analysed at T0 (data not shown). *MED12* cDNA sequencing demonstrated that three tumors had a point mutation in the codon 44 hotspot (c.131 G>A, p.G44D), while one tumor presented an indel mutation in exon 2 (c.118_124del7insC, p.N40_K42insQ) (Supplementary Figure S2). After LT-culture, the wild-type allele was not detected (L100) or barely detected (L94, L95, and L98), indicating that the mutation persisted throughout the entire culture period and the transcriptome profile mainly corresponds to *MED12* mutated SMCs (Supplementary Figure S2).

### Robust transcriptomic profiling enables characterization of long-term organ cultures

To characterize the transcriptome in the *ex vivo* culture slices, differential gene expression analyses were conducted on paired samples of UL and adjacent MM collected at baseline and after LT-culture. Following count normalization, both datasets demonstrated high-quality dispersion fits (Figures 1A and 1E). Principal component analysis (PCA) revealed distinct clustering between T0 and LT-culture in both tissue types (Figures 1B and 1F), with PC1 accounting for 70% and 77% of the total variance in UL and MM datasets, respectively. Tight intra-group clustering and the absence of outliers confirmed the consistency of samples across conditions. Volcano plots (Figures 1C and 1G) revealed extensive transcriptional remodelling, with 1,522 genes upregulated and 1,824 downregulated in the LT-culture of UL samples (Supplementary Dataset S1), and 1,958 upregulated and 2,322 downregulated genes in the LT-culture of MM tissues compared to T0 (Supplementary Dataset S2). Heatmaps of the most significantly differentially expressed genes (Figures 1D and 1H), filtered by Log2 FC > 1 or < -1 and q-value < 0.1, demonstrated clear condition-dependent clustering and consistent expression profiles across biological replicates. Together, these results indicate that LT-cultures induced robust and distinct transcriptomic patterns in both fibroid and myometrial tissues.

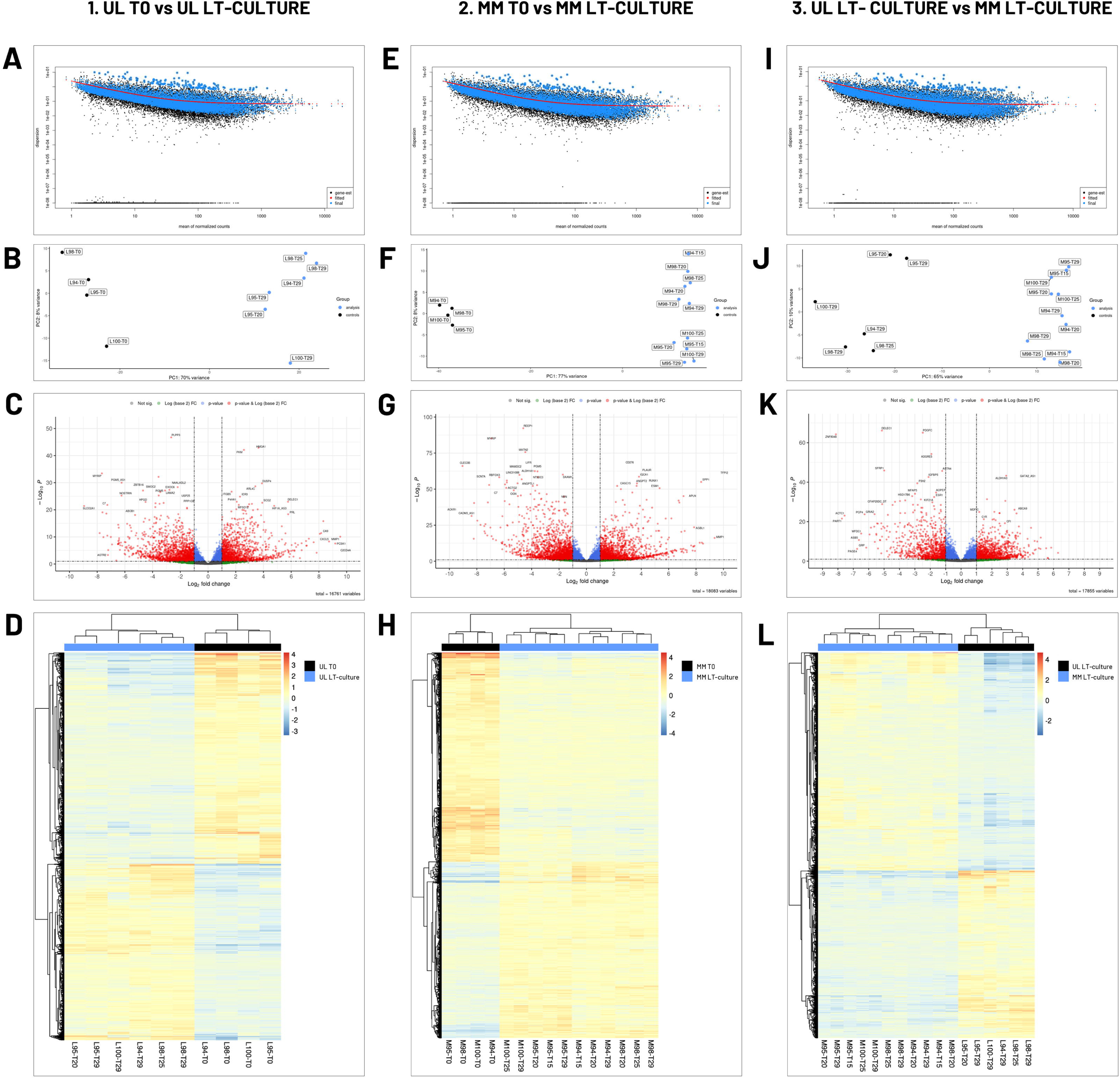

Differential gene expression analysis was also performed by comparing the LT-culture of fibroids with the LT-culture of matched myometrium. The dispersion model yielded a high-quality fit to the normalized count data (Figure 1I). PCA (Figure 1J) revealed clear segregation between UL LT-culture and MM LT-culture samples, with PC1 explaining 65% of the total variance. The volcano plot (Figure 1K) revealed extensive differential gene expression, identifying 1,444 significantly upregulated and 1,112 downregulated genes in the LT-culture of MM relative to the LT-culture of UL (Supplementary Dataset S3). Consistently, the heatmap of the most differentially expressed genes highlighted reproducible condition-specific expression patterns and clear sample clustering (Figure 1L).

### Hypoxia induced stem cell activation after long-term culture

Transcriptomic profiling revealed that in LT-culture, consistent activation of hypoxia-related pathways occurred in both UL and MM tissues when compared to their respective T0. In UL samples (Supplementary Figure S3), GO enrichment analysis identified a significant upregulation of pathways related to hypoxia, with the *response to oxygen levels* being among the top ten pathways with the most significant adjusted p-value (q = *8*.28E-06). Additionally, KEGG analysis revealed overrepresentation of the *HIF-1 signaling pathway* and *central carbon metabolism in cancer*, both indicative of metabolic adaptation to reduced oxygen availability (q = 1.05E-03 and 2.75E-05, respectively).

Similarly, GO pathways functionally aligned with responses to hypoxic stress were also prominently enriched in myometrial tissue (Supplementary Figure S4), such as *inflammatory response*, v*asculature development*, *tube morphogenesis*, *blood vessel development*, and *blood vessel morphogenesis*, all of which exhibited strong upregulation, with adjusted p-values ranging from 7.31E-08 to 3.81E-07. In parallel, KEGG analysis revealed significant enrichment of the *lysosome*, *HIF-1 signaling*, and *central carbon metabolism in cancer*, reinforcing the presence of hypoxia-driven metabolic reprogramming (q = 2.06E-07, 1.59E-03, and 2.37E-03, respectively). These findings collectively suggest that long-term organ cultures promote a hypoxia-like microenvironment within tissue slices, likely resulting from limited oxygen diffusion in the *ex vivo* model.

### Stem cell markers were differentially expressed in myometrium and leiomyoma long-term culture

Hypoxia is a widely accepted trigger of SC activation^16,17^. A transcriptomic comparison between tissues at T0 and those after LT-cultures revealed how hypoxia induced genetic reprogramming in normal and tumor SCs in their original niche (Supplementary Dataset S1 and S2). Substantial evidence supports the involvement of *HMGA* genes, which encode non-histone chromatin-associated proteins, such as *HMGA1* and *HMGA2* in promoting stem cell proliferation and stemness maintenance^23,24^. We detected both genes highly upregulated in normal and tumor cells after LT-cultures, particularly *HMGA1*, which was the most significantly upregulated gene in the UL LT-culture dataset (Table 1). Notably, *HMGA2* showed striking overexpression in myometrial tissues (FC = 46). *HMGA2* is an upstream regulator of the transcription factor *PLAG1*, frequently overexpressed in benign adipocytic tumors^25^. Accordingly, we observed significant upregulation of *PLAG1* in LT-culture of UL (FC = 5), but also in normal myometrium (FC = 4) (Supplementary Dataset S1 and S2).

**Table 1.** Differential Expression of Stemness and Differentiation-Related Genes in Long-Term Cultures (LT-Cultures). Genes shown were either overexpressed or downregulated in UL and MM LT-cultures compared to their respective T0 controls (no asterisk), in UL LT-cultures relative to MM LT-cultures (*), or in both comparisons (**). In the latter case, fold-change (FC) and adjusted p-value (q-value) correspond to the most statistically significant comparison.

| Upregulated Genes | UL FC | q-value | MM FC | q-value |
| --- | --- | --- | --- | --- |
| HMGA1 | 12.5 | 1.38E-43 | 12.8 | 6.32E-27 |
| HMGA2 | 16.5 | 3.22E-16 | 42.6 | 5.94E-11 |
| ITGA2 | 5.8 | 9.77E-08 | 5 | 3.23E-11 |
| ITGA2-AS1 |  |  | 10 | 0.019653 |
| ITG2-AS1 | 2 | 0,003023 |  |  |
| ITGA3 | 4.9 | 1,79E-12 |  |  |
| ITGA11 | 2.3 | 2,58E-05 |  |  |
| ITGAL |  |  | 2.5* | 0,096444 |
| ITGAX | 5.3 | 0,000143 | 9.5 | 1,67E-10 |
| ITGB1-DT |  |  | 2.6 | 0,015534 |
| ITGB2 |  |  | 2.6** | 2,92E-05 |
| ITGB2-AS1 |  |  | 6.8* | 0,04473 |
| ITGB5 | 3.2 | 2,03E-27 |  |  |
| ITGB7 | 6.8 | 0,040586 | 9.4** | 0,000128 |
| KLF3-AS1 | 2.2 | 0,001457 | 3.1 | 6,72E-06 |
| KLF5 | 5.4** | 5,27E-05 |  |  |
| KLF10 |  |  | 2.4 | 0.000348 |
| SOX4 |  |  | 6.3 | 2.39E-32 |
| SOX6 |  |  | 2.4* | 0.005727 |
| SOX7 |  |  | 9.8** | 1,47E-08 |
| SOX9 | 8.0 | 0,007551 | 17.5 | 0,010392 |
| SOX10 |  |  | 8.7* | 0,022779 |
| SOX11 | 3.2 | 0,050939 | 57.7 | 1,06E-06 |
| SOX17 |  |  | 4* | 0,002467 |
| SOX18 |  |  | 7.2** | 6,24E-07 |
| MYCN |  |  | 9.2 | 0.027 |
| MYCL | 4.8 | 0.048 |  |  |
| MYCT1 |  |  | 5.2* | 3,14E-05 |
| HOXA13 | 28.4* | 4E-16 |  |  |
| HOXA11-AS | 2.2* | 2,07E-06 |  |  |
| HOXA-AS2 |  |  | 4.2 | 0,020945 |
| HOXB2 |  |  | 2.0* | 0,005583 |
| HOXB3 |  |  | 4.0* | 2,08E-09 |
| HOXB5 |  |  | 9* | 2,9E-10 |
| HOXB6 |  |  | 7.2** | 1,18E-07 |
| HOXB7 |  |  | 10* | 8,35E-06 |
| HOXB8 |  |  | 27.5** | 2,49E-06 |
| HOXB9 |  |  | 3.8* | 0,068273 |
| HOXC4 |  |  | 2.5* | 0,046141 |
| HOXC8 |  |  | 6.5* | 0,004095 |
| HOXC9 |  |  | 4** | 0,002254 |
| HOXD8 |  |  | 2.3* | 0,010697 |
| HOTAIR |  |  | 10.2* | 0,006019 |
| CD24 | 9* | 4,52E-05 |  |  |
| CD34 |  |  | 5.5* | 0,001613 |
| CD73 (NT5E) | 2* | 9,83E-06 |  |  |
| CD90 (THY) | 2.5* | 8,1E-05 |  |  |
| KIT |  |  | 7.0** | 6,76E-13 |
| KITLG |  |  | 2.5* | 0,000561 |
| Downregulated Genes | UL FC | q-value | MM FC | q-value |
| ITGA6 | 3.4 | 1,53E-05 | 2.7 | 2,02E-07 |
| ITGA7 |  |  | 3.1 | 1,84E-05 |
| ITGA9 | 5.9 | 1,15E-05 | 3.8 | 3,84E-20 |
| ITGB4 | 78** | 1,25E-08 | 24 | 1,02E-23 |
| ITGB1BP2 |  |  | 10.3 | 3,64E-28 |
| ITGB8-AS1 | 9 | 0,013808 | 4.6 | 0,013204 |
| ITGBL1 | 4.2 | 0,006597 | 5.1 | 0,0001 |
| KLF2 | 8.1 | 9,13E-16 | 8.6 | 2,36E-14 |
| KLF2-DT |  |  | 3.6 | 0,021982 |
| KLF4 | 5 | 0,002279 | 7.1 | 4,33E-06 |
| KLF6 |  |  | 2 | 1,06E-09 |
| KLF9 |  |  | 2.5 | 3,7E-06 |
| KLF9-DT |  |  | 3.6 | 6,62E-08 |
| KLF11 | 2.2 | 0,068824 |  |  |
| KLF15 | 11.2 | 9,19E-05 | 9.5 | 2,69E-13 |
| KLF17 |  |  | 2.3 | 0,004381 |
| SOX6 | 3.5 | 2,83E-05 |  |  |
| SOX13 | 2.1 | 0,002733 |  |  |
| SOX2-OT |  |  | 4.0 | 2,42E-15 |
| HOXA13 |  |  | 4.5 | 0,036192 |
| HOXC4 |  |  | 3.3 | 0,001568 |
| HOXD-AS2 |  |  | 5.5 | 0,043619 |
| HOXAD8 | 2.2 | 0,033054 |  |  |

Similarly, the expression of integrin genes (*ITG*) is critically involved in both the proliferation and differentiation of stem cells. *ITGA2* (CD49b), previously identified in UL SCs, was also detected in MM LT-cultures (Table 1). Interestingly, MM cells exhibited a unique and marked overexpression of the long non-coding RNA *ITGA2-AS1* (FC = 10). Three integrins were specifically upregulated in UL LT-tissues: *ITGA3*, *ITGA11*, and *ITGB5*, the latter being the fifth most significantly upregulated gene in the UL LT-culture dataset (Supplementary Dataset S1, Table 1). Conversely, integrins *ITGAL* and *ITGB2* and the long-non coding RNAs (lncRNA) *ITGB1-DT* and *ITGB2-AS1* were uniquely upregulated in MM LT-culture (Table 1).

The Krüppel-like factors (*KLFs*) are pluripotency-associated transcription factors that play essential roles in stem cell biology^26^. *KLF5* and *KLF10* mRNAs were uniquely upregulated in UL and MM LT-culture, respectively (Table 1).

The SRY-related HMG-box (*SOX*) transcription factors play a critical role in maintaining stem cell identity and regulating lineage commitment^27^. Notably, six *SOX* genes were uniquely upregulated in MM long-term cultures, with four of them showing highly significant adjusted p-values (Table 1).

The *MYC* family of transcription factors is essential for cellular reprogramming and plays a pivotal role in regulating stem cell pluripotency and proliferation^28^. In this study, *MYCL* was upregulated in UL LT-cultures, whereas *MYCN* and *MYC target 1* (*MYCT1*) were specifically upregulated in MM LT-cultures (Table 1).

Homeobox (*HOX*) genes, known for their critical role in orchestrating embryonic development and tissue differentiation, exhibited distinct expression profiles between UL and MM LT-cultures. Only *HOXA13* and *HOXA11-AS* were significantly upregulated in UL LT-culture (Table 1). In contrast, a wide array of *HOX* genes, including members of the *HOXB*, *HOXC*, and *HOXD* clusters, along with the lncRNA *HOTAIR*, were markedly upregulated in MM LT-culture.

Classical SC markers already found upregulated in UL were also overexpressed in our dataset, including *CD24*, *CD73*/*NT5E*, and *CD90*/*THY1*, whereas *CD34* was upregulated in MM LT-culture. Interestingly, *KIT* uniquely increased expression in MM LT-culture slices, like *KITLG* ligand (Table 1).

While activation of SC genes maintains pluripotency, their downregulation enhances differentiation toward specific lineages. *ITGA7* and *ITGB1BP2* decreased expression uniquely in MM LT-cultures (Table 1). *KLF2* and *KLF15*, both known to play roles in maintaining stem-like characteristics, were downregulated in both UL and MM LT-cultures with highly significant q-values. *KLF4*, a key pluripotency-associated factor, was similarly decreased in both tissues. Additional *KLF* members such as *KLF6*, *KLF9*, and *KLF17*, along with two lncRNA divergent transcripts (*KLF2-DT, KLF9-DT*), were selectively inhibited in MM, whereas *KLF11* was uniquely downregulated in UL LT-culture. *SOX* 6 and *SOX13* were uniquely downregulated in UL, while *SOX2-OT*, a lncRNA that promotes *SOX2* expression, was downregulated in MM LT-cultures^29^. Together, these findings highlight a dual transcriptional program induced in LT-culture under hypoxic conditions: while key stemness genes were differentially upregulated in UL and MM slices, there was a concurrent downregulation of genes associated with pluripotency maintenance, suggesting a coordinated transition toward lineage differentiation that differs between fibroid and myometrial tissues.

### Differential metabolic gene activation in myometrium and leiomyoma long-term culture

Reactome enrichment pathways analysis revealed that cells in LT-culture activated central metabolic pathways in both UL and MM, aligning with the energetic and biosynthetic demands of SC activation and differentiation (Supplementary Figure S5), including *Metabolism of Carbohydrates*, *Glucose Metabolism*, *Glycolysis*, and *Gluconeogenesis*. However, tissue-specific gene analysis revealed distinct transcriptional signatures (Supplementary Dataset S1 and S2). Genes upregulated in glycolysis and glycogen metabolism in UL LT-cultures (*GAPDH*, *PCK1*, *PGM1*, *HKDC1*, *SORD*, *GBE1*, and *GYS1*) were different from those found in MM LT-cultures (*HK3*, *PC*, and *PCK2*). While UL uniquely activated genes related to the degradation of complex carbohydrates (*MANBA* and *HYAL3*), MM activated genes involved in sugar transport or processing (*AKR1B1*, *SLC37A2, SLC37A4*, and *SLC35B2*). Moreover, pathways related to SLC-mediated transmembrane transport for bile salts, organic acids, metal ions, and amine compounds were activated only in fibroids LT-cultures (Figure 2A).

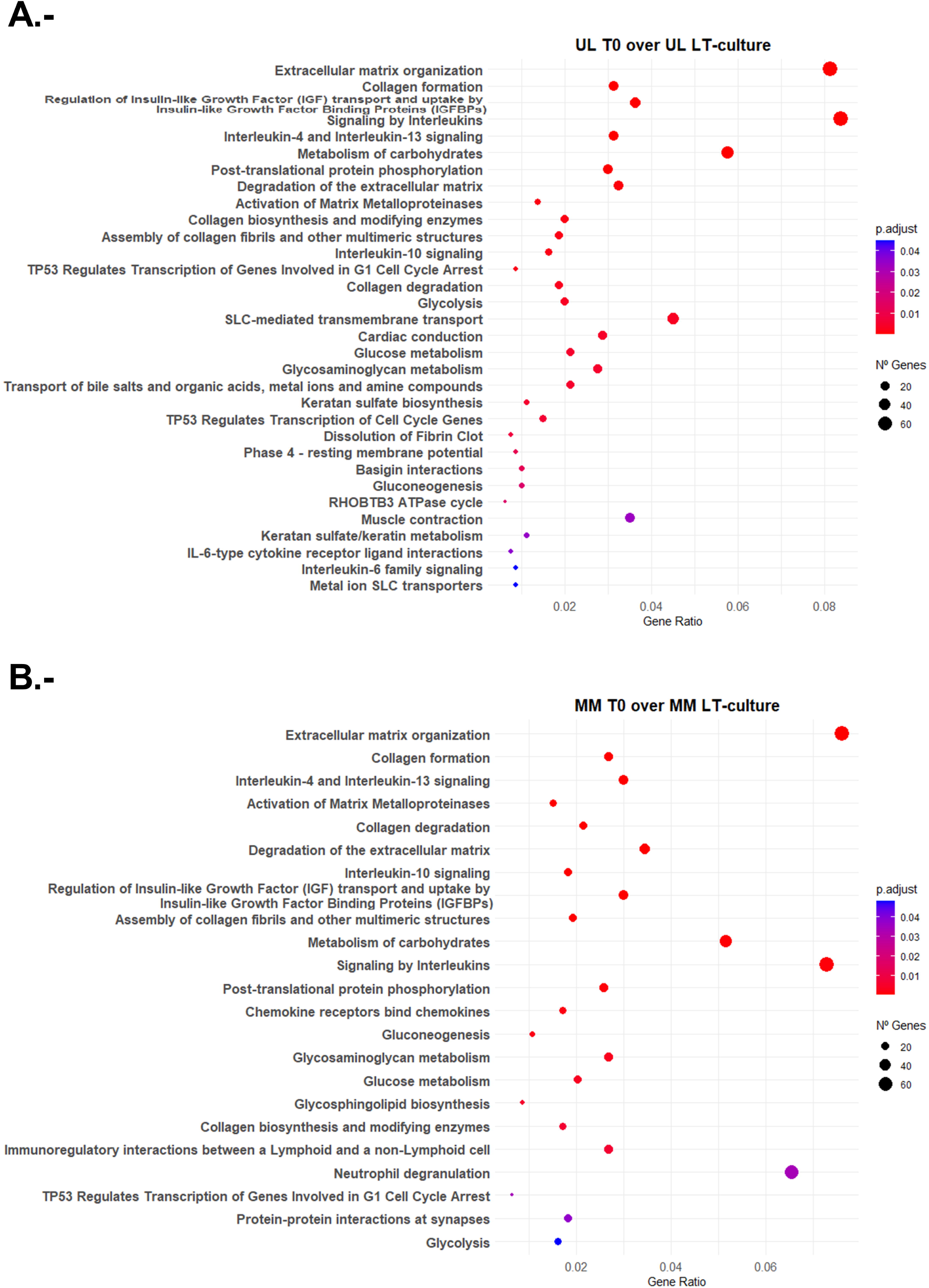

Interestingly, a previous study found that *PLIN2*, a lipid droplet-binding protein found on the surface of lipid droplets in most mammalian cell types, was downregulated in UL. Loss of *PLIN2* enhances both mitochondrial activity and glycolytic flux, driving a metabolic reprogramming that promotes a more proliferative cellular phenotype in UL cells^30^. Accordingly, we found upregulated expression of *PLIN2* in MM LT-culture compared to UL LT-culture (FC = 3.2, q = 1.49E-10) (Supplementary Dataset S3). Overall, these findings underscore a shared metabolic reprogramming during LT-culture in normal and tumor tissue, while also revealing distinct, tissue-specific metabolic adaptations during SC proliferation and differentiation.

### Long-term culture uncovers differential ECM remodelling in myometrium and leiomyoma

A defining hallmark of uterine fibroids is their abundant and disorganized ECM, which contributes to the increased tissue stiffness and the characteristic fibrotic appearance of this tumor^31^. UL and MM LT-cultures exhibited extensive matrix remodelling, with eight core ECM-related pathways commonly upregulated in both tissues when comparing T0 samples to LT-cultures (Supplementary Figure S5). In addition, UL showed exclusive activation of additional ECM-related pathways, such as *Basigin interactions*, *Keratan sulfate/keratin metabolism and biosynthesis*, and *Dissolution of fibrin clot* (Figure 2A). Thus, UL showed strong upregulation of pathways linked to proteoglycan synthesis and modification (*B4GALT4, B4GALT5, CHST1, CHST6, ST3GAL1, ST3GAL2, ACAN, FMOD*), adhesion and matrix remodelling (*ITGA3, MMP1*), and fibrinolysis (*PLAT, PLAU, PLAUR, SERPINE1, SERPINE2*). Conversely, MM uniquely overexpressed two pathways: *Glycosphingolipid biosynthesis* (*B3GNT5, B4GALNT1, CERK, ST3GAL2, UGCG, UGT8*), and *Neutrophil degranulation* (*MMP8, MMP9, PLAU, PLAUR, S100A8, S100A9, ITGAX, ITGB2*), consistent with immune-mediated ECM remodelling (Figure 2B). Notably, when comparing UL LT-cultures to MM LT-cultures, three ECM pathways were significantly downregulated in MM, indicating their preferential upregulation in UL (Figure 3), including *Defective B3GALTL causes PpS*, *O-glycosylation of TSR domain-containing proteins*, and *Diseases of glycosylation*. Gene pathways analysis revealed UL enrichment for ECM crosslinking enzymes and glycoproteins (*ADAMTS17, ADAMTS19, ADAMTS2, ADAMTS8, THBS1, THSD4*), structural proteoglycans (*BCAN, CSPG4, FMOD, OGN, PRELP, SDC3*), and glycosylation machinery (*DPM2, MUC12, SBSPON, SEMA5A*). Overall, these findings highlight a profound divergence in extracellular matrix dynamics between UL and MM proliferation.

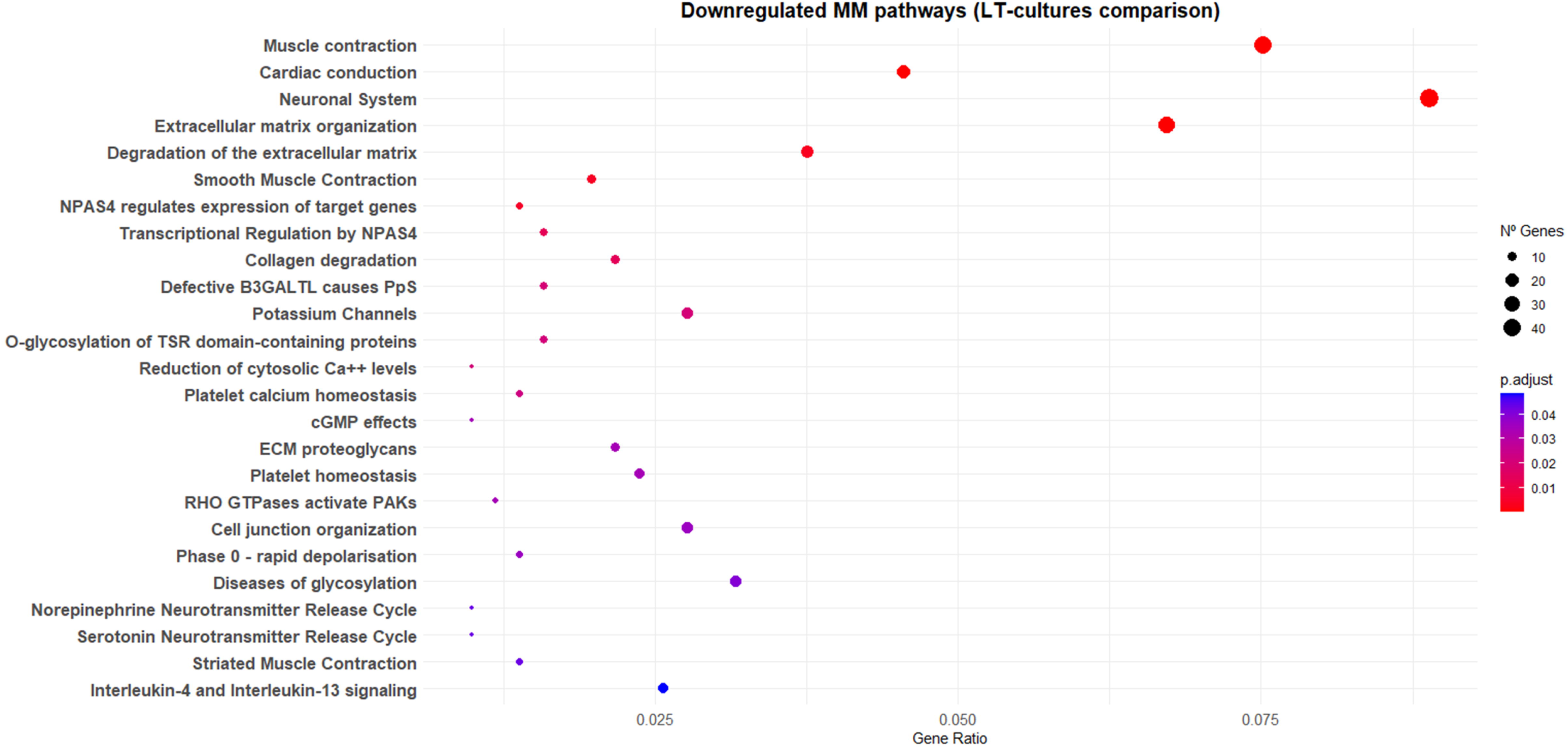

### Immune surveillance and angiogenesis in the myometrium are absent in leiomyoma long-term culture

Transcriptomic analysis comparing UL LT-cultures to MM-LT cultures revealed a striking enrichment of immune and vascular-related pathways in MM (Figure 4). Upregulated pathways included both innate and adaptive immune processes, as well as vascular homeostasis and repair-associated pathways. Conversely, UL-LT cultures showed upregulation of the *Interleukin-4 and Interleukin-13 signalling* pathway (Figure 3), whereas comparison of UL-T0 to UL-LT revealed enrichment of IL-6-type cytokine signalling (Figure 2A). Interestingly, the Phospholipid Phosphatase 3 (*PLPP3*) was the most significantly downregulated gene of the UL dataset (FC = 6.3, q = 1.855E-47, Supplementary Dataset S1). *PLPP3* regulates vascular homeostasis by dephosphorylating lipid mediators like lysophosphatidic acid and sphingosine-1-phosphate, thereby controlling endothelial cell adhesion, migration, and inflammatory signalling^32^.

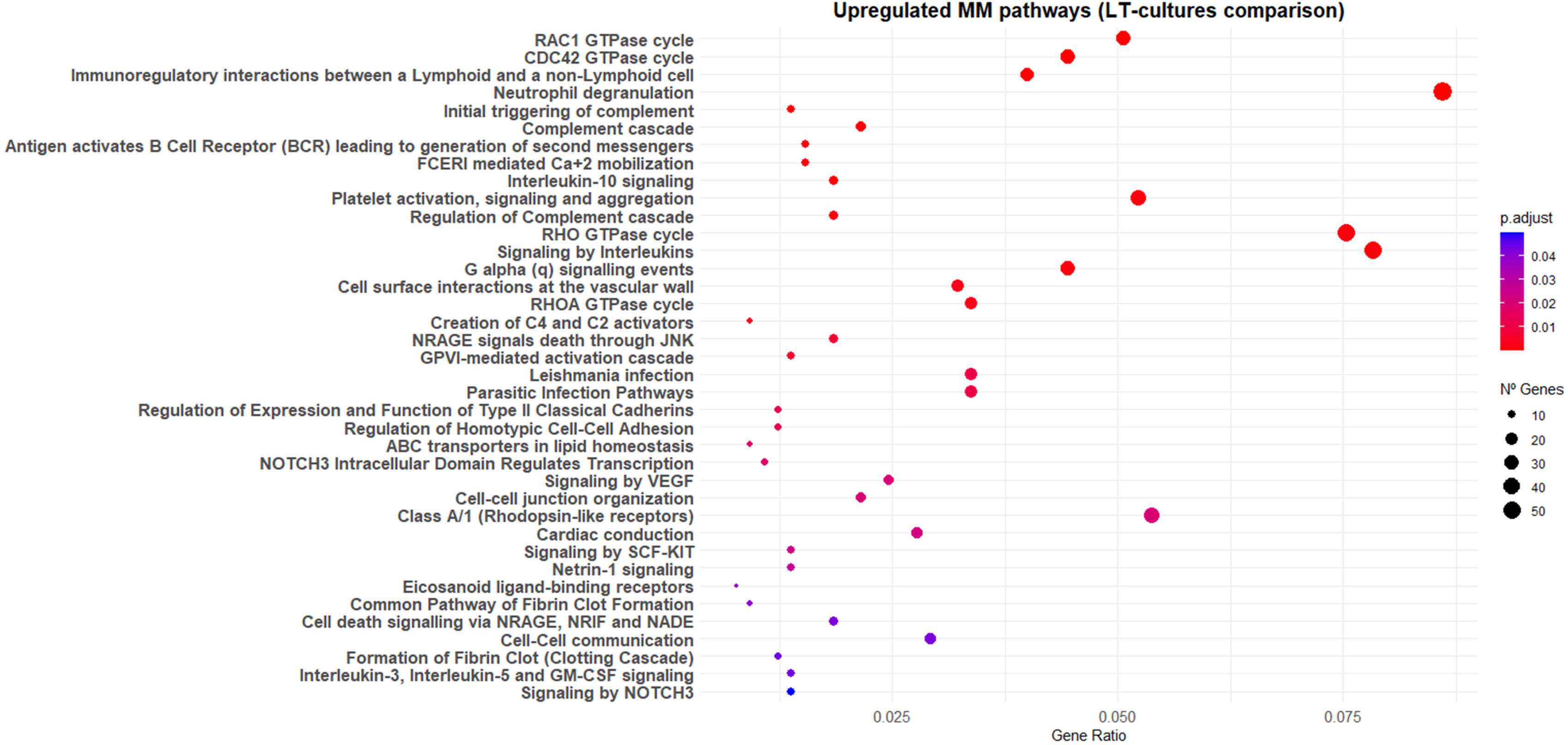

Overall, these patterns suggest robust immune surveillance and active vascular maintenance in MM, potentially aimed at preserving tissue integrity and promoting repair. Conversely, UL exhibited a more restricted, Th2-biased immune profile typically associated with fibrosis and immune evasion^33^.

### Fibroids exhibit a self-sustained secretory-contractile program in long-term culture

Transcriptomic comparison between LT-cultured normal and tumor tissues revealed a significant downregulation in MM of pathways related to neurotransmitter release, the neuronal system and transcriptional regulation by *NPAS4* (Figure 3). These genes (such as *SYT1, SNAP25, RIMS1, SYN3*, and the neuronal activity-dependent transcription factor *NPAS4*) are essential for vesicle trafficking and neurotransmitter release in the central nervous system (CNS). Notably, NPAS4 has been shown to play a significant role in vesicle dynamics and optimizing cellular metabolism during excitation in neuroendocrine cells^34^. These genes are not exclusive to the CNS and they show detectable (low, moderate or high) RNA expression in cardiac, skeletal, and smooth muscle cells (The Human Protein Atlas, 2025)^35^. The upregulation of these vesicle-related genes in UL (Supplementary Dataset S3) suggests that fibroid smooth muscle cells may acquire a more secretory phenotype during proliferation.

The transcriptomic profile of UL LT-culture compared to MM LT-culture suggests an enhanced ability for self-directed contractile engagement in UL, as evidenced by the coordinated overexpression of pathways related to muscle contraction and enhanced electrical excitability (Figure 3). The downregulation of *Phase 0 – Rapid Depolarization and Potassium Channels* in MM over UL suggests that fibroid smooth muscle cells can generate or respond to action potentials without the need for intense external stimulation. At the same time, the *Reduction of Cytosolic Ca²⁺ Levels* pathway overexpressed in UL suggests that SMCs can rapidly return to their basal state after activation. Moreover, the unique activation of multiple pathways related to muscle contraction (both smooth and striated) suggests that fibroid myocytes’ contractile machinery is readily activatable (Figure 3). Conversely, different pathways regulating contractility, cytoskeletal dynamics, and calcium signalling were activated in MM (Figure 4), including RAC1, CDC42, RHO GTPase cycles, the high-affinity IgE receptors (FCεRI) pathway, typically associated with the immune system triggering the release of Ca²⁺ from intracellular stores (*FCERI Mediated Ca²⁺ Mobilization*), and pathways related to receptors responsive to hormones, prostaglandins, and leukotrienes involved in contraction (*G alpha (q) signaling events, Rhodopsin-like Receptors, and Eicosanoid Ligand-Binding Receptors*).

Overall, these data suggest a divergence from the tightly regulated, signal-dependent and immune-modulated contractile apparatus in MM from a more self-sustained secretory-contractile mechanism in UL.

### Myometrium activates cell-death mechanisms absent in fibroids during long-term culture

After the repopulation phase that typically begins between days 15 and 29, cells in the tissue slices finally disappear. However, the mechanism of cell death differs in fibroids and healthy myometrium. Comparison of LT-culture of UL over LT-cultures of MM revealed consistent upregulation of apoptosis-related pathways in MM, such as *RAGE signals death through JNK, signalling by NOTCH3*, that influences transcription of genes involved in cell cycle arrest and apoptosis, and *Cell death signaling via NRAGE, NRIF, and NADE* (Figure 4). In contrast, fibroids show an absence of apoptotic signalling, while the RNA component of telomerase (*TERC*) was upregulated (FC = 5.3, q = 4.64E-06, Supplementary Dataset S3). This data suggests that myometrial SMCs are actively engaging programmed cell death mechanisms that are not present in UL, which is more prone to regulating cellular longevity.

### Commonly deregulated pathways in fibroids were also detected in long-term leiomyoma organ cultures

In UL LT-cultures, we found several signalling pathways that have been consistently reported as dysregulated in fibroids, either compared to their respective T0 controls (no asterisk), in UL LT-cultures relative to MM LT-cultures (*), or in both comparisons (**) (Supplementary Datasets S3, S4 and S5). Reactome pathway enrichment analysis revealed that the *Regulation of Insulin-like Growth Factor (IGF) transport and uptake by Insulin-like Growth Factor Binding Proteins (IGFBPs)* pathway was upregulated in both UL and MM LT-cultures (Supplementary Figure S5, Figure 2). However, important differences were observed analysing individual members of the IGF family. Thus, *IGF1* (FC = 4.4, q = 1.83042E-13), *IGF1R\** (FC = 2.5, q = 1.03827E-18), *IGF2R* (FC = 2.6, q = 2.98168E-15), *IGFBP5\*\** (FC = 4.3, q = 2.481E-43), *IGF2BP2* (FC = 4, q = 0.001827543), *IGFBP7-AS1\** (FC = 3.5, q = 0.000699113), *IGFBPL1\** (FC = 3.8, q= 0.038506339), *IGFL2\** (FC = 14, q= 2.16069E-13), *IGFL3\*\** (FC = 36, q = 4.1194E-17), *IGFL4\** (FC = 15, q = 0,004079777), *IGFN1* (FC = 10.6, q = 1.06827E-08) were uniquely upregulated in UL, whereas *IGFBP1\*\** (FC = 48, q = 0.023415129, *IGFBP3* (FC = 7.5, q=0.000124338) and *IGFBP6\** (FC = 2.4, q= 0.001350797) increased expression only in MM LT-cultures.

The Wnt signalling pathway has been strongly implicated in leiomyoma formation^36^. In UL LT-cultures, we observed upregulation of several Wnt pathway components. Notably, *WNT3\** was significantly upregulated (FC = 3.4; q = 0.025112), with potential to interact with frizzled class receptors to mediate signal transduction to the nucleus, such as *FZD1\** and *FZD7\**, which were also upregulated in UL (FC = 2.2; q = 8.78E-6 and FC = 3.2; q = 2.42E-9, respectively). Consistently, we observed elevated expression of cyclin D1 (*CCND1\**) (FC = 2.2; q = 0.001442), a well-established downstream target of Wnt. Interestingly, in response to hypoxia, beta-catenin increased mRNA stabilization of the stem cell regulator *SNAI2* ^37^, which we found highly significantly upregulated in UL LT-culture (*SNAI2*,* FC = 4.0, q = 2.7E-11). Finally, secreted frizzled-related protein 1 (*SFRP1\**, FC = 31; q = 1.19E-46) and *NKD1\** (FC = 15, q = 3.51E-20) were among the most significantly upregulated genes in our dataset, corroborating previous findings^5,36^.

The family of transforming growth factor beta ligands have been implicated in the development and progression of UL. Several studies have found that *TFGB1*, *TFGB2*, and *TFGB3* were upregulated in UL compared to matched MM^38–42^. Similarly, we found significant upregulation of *TFGB1* (FC = 2.5, q = 0.000547), *TFGB2*\* (FC = 6.0, q = 1.21E-25), and *TFGB3*\* (FC = 2.2, q = 2.41E-06), as well as the transforming growth factor beta-induced *TGFBI* (FC = 5.6, q = 1.72E-12). Interestingly, in MM LT-cultures over T0 we observed downregulation of transforming growth factor beta 1 induced transcript 1 (*TGFB1I1*, FC = -2.1, q = 2.97E-06) and the TGF-beta receptors *TGFBR1* (FC = -2.2 q = 1.13E-07), *TGFBR2* (FC = -3.1, q = 2.87E-36), and *TGFBR3* (FC = -5.6, q= 2.14E-22). Activin-A, a member of the TGF-β superfamily, is a protein formed by the dimerization of two beta-A subunits encoded by the *INHBA* gene. Increased levels of activin-A mRNA have been detected in UL, where it increased the expression of several ECM components *in vitro*, suggesting a profibrotic role in UL development^43^. In our dataset, *INHBA*\*\* was found to be significantly upregulated (FC = 3.7, q = 3.8E-06).

There is strong evidence that aberrations in the Retinoid Acid-signalling pathway play a role in leiomyoma development and growth^44–48^. While *ADH1* and *ALDH1* were found to be downregulated, the cellular retinoic acid binding protein 2 (*CRABP2*) expression was upregulated. We detected *ADH1B*\*\* (FC = -121, q = 6.17E-08), *ALDH1A1*\*\* (FC = -14, q = 5.12E-07), *ALDH1A3*\*\* (FC = -7.5, q = 5.02E-44), *ALDH5A1* (FC = -3.0 q = 0.005217) *ALDH6A1* (FC = -3.2 q = 1.49E-08), *ALDH7A1* (FC = -2.7 q = 3.31E-12) downregulated while *CRABP2*\* (FC = 4.8 q = 6.64E-11) increased expression in UL LT-cultures.

UL is considered highly responsive to ovarian steroid hormones due to the increased expression of sex steroid receptors. Additionally, aromatase expression by tumor cells is thought to play a key role in the growth and maintenance of leiomyomas. In agreement with these findings, we observed upregulation of estrogen receptor 1 (*ESR1\**, FC = 3.4, q = 3.22E-33), progesterone receptor (*PGR\**, FC = 3.0, q = 1.28E-10), and aromatase (*CYP19A1\**, FC = 7.7, q = 0.003167) when comparing MM and UL LT-cultures.

An overwhelming body of evidence linked increased expression of growth factors, particularly platelet derivate growth factors (PDGF)^31,49–54^ and fibroblast growth factors (FGF) and their two receptors, FGFR1 and FGFR2^55–58^ to enhanced cell proliferation and ECM accumulation in fibroids. In our dataset, *PDGFC\** was one of the five most significantly upregulated genes in UL LT-cultures (FC = 5.6, q = 8.49E-66), while *PDGFA* (FC = -2.5, q = 0.000971) and *PDGFD* (FC = -4.7, q = 1.27E-12) were repressed in MM T0 over MM LT-cultures. In addition, *FGF14\** (FC = 3.0, q = 3.05E-12) and the two receptors, *FGFR1\** (FC = 2.0, q = 2.05E-12) and *FGFR2\** (FC = 2.5, q = 0.000153), were also upregulated in UL LT-culture.

Recently, high upregulation of the tryptophan (Trp) 2,3-dioxygenase (*TDO2*) gene has been detected in *MED12-*mutated UL compared to matched myometrium, with functional studies indicating an essential role in tumor growth^59–61^. Consistent with these reports, we found a strong upregulation of *TDO2* mRNA (*TDO2*,* FC = 14.8, q = 2.51E-05) in UL LT-cultures.

Overall, these findings suggest that LT-cultures of fibroids faithfully recapitulate the characteristics of the original tumors, providing a robust and unique experimental platform for studying the mechanisms underlying SC activation and growth.

## Discussion

In this study, we demonstrate for the first time that long-term organ cultures of myometrial and fibroid tissue slices spontaneously develop hypoxic conditions (Supplementary Figures S3 and S4). Due to the considerable thickness of the tissue slices (around 500 μm), oxygen diffusion is limited, leading to a gradual widespread cell death throughout most of the slice starting from day seven of culture. Remarkably, quiescent somatic SC remained viable and become hypoxia-activated in their niche around 15 days of the organ culture, resulting in a unique biological scenario where SCs proliferate and differentiate into myocytes in the native ECM. Comparative transcriptomic analysis between freshly collected tissue and LT-culture reveals the transcriptional reprogramming driven by normal and tumor SC activation, proliferation, and differentiation. Furthermore, comparison of normal and tumor tissue at LT-culture reveals additional differences in gene expression patterns.

Immunohistochemical analysis in LT-cultures mainly detected DES+/VIM+ cells, corresponding to SMC in both MM and UL cultures (Supplementary Figure S1). Single-cell sequencing analyses revealed three distinct SMC clusters in UL and MM cells, with the SMC cluster 0 characterized by expression of *TAGLN*, *CNN1*, *ACTA2*, and *CACNA1C*^62^. In addition, spatial transcriptomic analysis of fibroid tissue detected expression of *ACTG2* as specific markers of SMC^63^. We found all these SMC markers highly significantly upregulated in UL LT-culture compared to MM LT-culture (q > 10-7, supplementary Dataset S3). These genes encode proteins central to smooth muscle contraction and cytoskeletal integrity, supporting a myocyte repopulation with a more contractile phenotype in UL, as evidenced by the reactome enrichment analysis (Figure 3). Importantly, the *MED12* mutant allele found at T0 in the four UL remained preferentially expressed in the LT-cultures, supporting the hypothesis that fibroids arise from a mutated somatic SC^12,13^.

Nevertheless, other known cell types present in UL and MM tissues may also exist in LT-cultures. We detected a highly significant upregulation of several fibroblast-related genes in UL-LT over T0 (Supplementary Dataset S1), including *SERPINE1*, *SERPINE2*, *IER2* and *NR4A2*^62^. *SERPINE1* is notably enriched in cancer-associated fibroblasts, where it promotes extracellular matrix remodeling, cell migration, and tissue fibrosis. Similarly, *SERPINE2*, *IER2*, and *NR4A2* contribute to fibroblast-driven processes such as collagen synthesis, motility, and inflammatory signalling^64–67^. In addition, a comparison between UL and MM LT-culture showed typical endothelial cell (EC) markers, such as *PECAM1* (*CD31*), *VWF*, and *CDH5* (*CD144*), upregulated in MM LT-culture (Supplementary Dataset S3). Up to eight EC clusters have been detected in both MM and UL^62^. On the other hand, markers of immune cells such a monocyte/macrophage (*CD14, CD163, CD163L1, CD68, CD36, CD300A, CD300LB, CD33, TPSB2*), lymphocytes T, B, NK and dendritic cells (*CD4, CD22, CD28, CD38, CD40, CD48, CD52, CD74, CD84*) were augmented in MM LT-culture when compared to UL LT-culture (Supplementary Dataset S3), suggesting residual immune cells preferentially present in MM slices. Immune cells have been found in both MM and UL tissue with a particular cell repertoire and cytokines profile expression^68–71^. Notably, the upregulation of typical markers of endothelial and immune cells and the significant enrichment of pathways related to vascular remodelling and innate and acquired immunity in MM LT-culture suggest that myometrial growth is accompanied by enhancing tissue vascularization, immune-mediated vascular remodelling, and immune surveillance during proliferation. Conversely, the increased expression of fibroblast-related markers, decreased angiogenesis, and cells of the immune system favour UL cell proliferation characterized by fibrotic ECM remodelling, immune evasion and abnormal angiogenesis. Accordingly, tumor-associated fibroblasts were particularly abundant in *MED12* mutated UL, with paracrine interactions between both cell types probably favouring fibroid growth^72^. Deficient and dysregulated angiogenesis was found in UL, with more expression of anti-angiogenic genes and less expression of pro-angiogenic ones^73,74^. Interestingly, we found *CD274* (*PD-L1*), an immune inhibitory receptor ligand that enables tumor cells to evade the immune system and promotes tumor growth^75^, significantly upregulated in UL LT-cultures (FC = 3.0, q = 0.015825, Supplementary Dataset S3). Recent data suggest that fibroid growth is associated with an immunosuppressive phenotype that facilitates ECM deposition^70^.

Functionally, the secretory phenotype detected in UL aligns with the abundant endoplasmic reticulum observed in fibroid myocytes, suggesting an increased synthesis capacity^76^, while it retained an readily activatable contractility, facilitating autonomous contraction. Conversely, MM myocytes exhibit contractility that is largely modulated by immune, hormonal, and paracrine cues, reflecting a more adaptive and finely regulated functional state. Notably, MM myocytes in long-term culture engage apoptotic pathways in response to prolonged stress, including RAGE-JNK and NOTCH3-mediated signals, indicating a regulated mechanism of programmed cell death. UL myocytes, by contrast, show an absence of apoptotic signalling, suggesting intrinsic resistance to programmed cell death. Ultrastructural analysis further revealed phagosomes within UL myocytes, consistent with autophagocytosis as the primary route for cellular clearance of UL SMC cells^76^. This intrinsic resistance is supported by previous studies demonstrating that in UL, elevated expression of anti-apoptotic proteins such as Bcl-2, diminished caspase-3 activity, and phosphoproteomic profiles favor cell survival over death^77,78^.

Several genes related to stemness maintenance and proliferation were found dysregulated in LT-culture, such as *HMG, ITG, KLF, SOX,* and *HOX* genes. *HMGA1* and *HMGA2* encode for non-histone DNA-binding proteins that regulate transcription by influencing DNA conformation. Both proteins are highly expressed during embryogenesis, but their expression is repressed or nearly absent in most adult tissues, except in some somatic SC populations, supporting their self-renewal and pluripotency^23,79^. Interestingly, during skeletal muscle differentiation, *HMGA2* mRNA levels increased after stem cell activation, remained upregulated during myoblast proliferation, and gradually decreased when myoblasts were induced to differentiate^80^. We detected both genes highly significantly upregulated in MM LT-culture, suggesting a fundamental role in myometrium SMC proliferation and uterine physiology. In fibroids, none of the tumors at T0 showed *HMGA2* amplification, indicating that the *MED12* mutation was the dominant driver alteration. However, in our dataset, the four fibroids with *MED12* mutations also exhibited a highly significant increase in *HMGA1* and *HMGA2* expression in LT-culture. Consistent with our findings, George et al.^81^ reported a subset of *MED12*-mutant fibroids exhibiting elevated *HMGA2* expression. In addition, the transcriptional profile of fibroid subtypes demonstrated that the *MED12* and *HMGA2* types shared the most differentially expressed genes and commonly dysregulated pathways^5^. We observed that *HMGA1* and *HMGA2* overexpression was detected between T0 and LT-culture in both MM and UL. Still, no deregulated expression of *HMGA1* or *HMGA2* was observed when directly comparing MM LT-with UL LT-culture. These findings suggest that in comparative analyses between fibroids and myometrium, *HMGA2* expression differences may not be apparent if the myometrial samples are in a proliferative state with elevated *HMGA2* expression, such as during the secretory phase, or the adjacent MM of patient is in an early stage of transformation with already molecular and structural changes that may predispose to fibroid development^82,83^. Interestingly, a previous study demonstrated that *HMGA2* upregulation is not only a common phenomenon in most fibroids, but some matched myometrium also showed augmented expression^84^.

Integrin receptors are the main molecular link between cells and the ECM, as well as cell–cell interactions. Due to the dynamic interactions with the ECM, ITGs play pivotal roles in regulating cell proliferation, differentiation, and migration^85^. The integrin ITGA2 has previously been associated with stemness and the initiation of metastasis in prostate and breast tumor cells, whereas an antibody against this marker successfully isolated UL SC^14,86^. We observed *ITGA2* overexpression in LT-cultures from tumor and normal cells. However, MM cells uniquely expressed the long non-coding RNA (lncRNA) *ITGA2-AS1* that negatively regulates *ITGA2* gene expression^86^. Indeed, the proliferative and metastatic effects induced by ITGA2 are mediated by the metabolic gene *ACLY*^86^, and we observed *ACLY* upregulation exclusively in LT UL-cultures (FC = 2.4, q = 2.29E-8). The more regulated expression of *ITAG2* in MM is supported by our previous immunohistochemistry analysis, where we detected few ITGA2-positive (CD49b) cells in MM LT-culture compared to large clusters of positive cells in UL LT-culture (Supplementary Figure S6).

Additionally, UL LT-cultures overexpressed *ITGA3*, *ITGA11*, and *ITGB5* genes implicated in tumor progression, metastasis, and poor survival across several cancer types^87–91^. Conversely, *ITGB2*, *ITGB2-AS1* and *ITGAL* were upregulated in MM LT-cultures (Table 1). ITGB2 suppresses tumor growth and metastasis in various cancers, whereas *ITGB2-AS1* lncRNA appears to enhance *ITGB2* expression^92,93^. ITGAL together with ITGB2 forms the αLβ2 integrin, which is expressed on leukocytes.

ITGAL facilitates immune cell adhesion and trafficking by binding to intercellular adhesion molecules, promotes T cell activation, enhances NK and cytotoxic T cell killing, and contributes to macrophage-mediated clearance of apoptotic cells, particularly neutrophils^94–96^. *ITGA7* was downregulated in MM LT-culture, and depending on the tumor context, it may function either as a tumor suppressor or oncogene. In glioblastoma and astrocytoma, endometrial, hepatocellular, and non-small cell lung cancers, it acts as an oncogene that promotes proliferation, stemness, invasion, and poor prognosis^97–100^. Together, we observed a tumor-specific upregulation of integrins associated with proliferation and metastasis in UL, while MM exhibits a more immunomodulatory integrin signature.

The *KLF* family of transcription factors play a crucial role in regulating stemness and maintaining pluripotency in SCs. Overexpression of *KLF4* enhances self-renewal, while its depletion leads to differentiation^101^. The downregulation of *KLF4* in UL and MM LT-cultures suggests an active differentiation process in cultured slices. Interestingly, *KLF5*, which was found upregulated in colorectal, cervical, breast and endometrial cancer, where it may act as an oncogene, was significantly overexpressed in UL LT-culture^102–105^. Conversely, *KLF10*, upregulated in MM LT-culture (Table 1), is a TGF-β–responsive transcription factor with a central role in fibrosis and muscle regulation.

KLF10 inhibits TGF-β–mediated stellate cell activation and fibrogenesis in the liver, whereas loss of KLF10 increases fibrosis and impairs skeletal muscle function^106–108^. The lack of KLF10’s inhibitory function appears to correlate with the elevated expression of TGF-β–related ECM genes in UL LT-culture, including *TGFB2*, *TGFB3, SCUBE3, EMILIN1* and *THBS1* (Supplementary Dataset S1 and S3). *KLF6* acts as a tumor suppressor gene in many cancers, but its alternative splice variant, *KLF6-SV1*, can promote tumor growth and metastasis^109–111^. This splice variant was upregulated in uterine leiomyosarcoma (LMS), where it correlates with increased proliferation, invasion, and resistance to apoptosis^112^. Transgenic mice expressing *KLF6-SV1* spontaneously develop LMS-like tumors, whereas silencing *KLF6-SV1* in LMS cell lines led to cell cycle arrest and enhanced apoptosis^112^. Interestingly, in our dataset, *KLF6* was significantly downregulated in MM LT-cultures (Supplementary Dataset S2). However, both transcripts have the same 3’ mRNA sequence, and the 3**’** RNAseq strategy cannot distinguish whether this corresponds to the canonical *KLF6* transcript or the oncogenic *KLF6-SV1* splice variant. Further studies are necessary to elucidate the role of KLF6 in fibroid proliferation, and particularly in those fibroids that can evolve to LMS. Other differentially expressed KLF genes included *KLF9, FLF11*, *FLF17* and the divergent lnc-RNA transcripts *KLF9-DT*. All of them have been involved in cancer development and progression^113^. Particularly, *KLF11* may act as tumor suppressor, inhibiting neoplastic transformation and cell proliferation, and it is frequently downregulated in various cancers^114^. Previous studies found reduced expression in uterine leiomyomas compared to matched normal myometrium, and this repression has been linked to epigenetic silencing via promoter hypermethylation^115,116^. Accordingly, we find significant downregulation of *KLF11* in UL LT-culture (Table 1). Overall, the expression patterns of *KLF* transcription factors suggest that in UL LT-cultures, pro-fibrotic and oncogenic pathways are activated, while in MM LT-cultures, cells maintain tumor-suppressor gene expression.

The *SOX* transcription factors are master regulatory genes controlling development and are fundamental to the establishment of sex determination in a multitude of organisms^117^. Several *SOX* genes were uniquely upregulated in MM LT-culture (Table 1). SOX4 is involved in uterine development and function, and its expression is tightly regulated by ovarian hormones^118^. Moreover, SOX4 is also expressed in the myometrium during labour, where it could play a crucial role in modulating the phenotypic switch of SMCs from relative quiescence to contractility^119^. SOX6 has been described as a tumor-suppressor gene in several cancers, suppressing cell proliferation by stabilizing p53 protein and subsequently upregulating p21^120^. SOX18 plays a vital role during embryogenesis by regulating blood vessel formation, guiding lymphatic differentiation, and determining endothelial cell fate^121^. Conversely, SOX13, downregulated in UL LT-cultures, may promote endothelial quiescence by suppressing inflammatory chemokines, helping preserve vascular integrity and immune homeostasis^122^. SOX2 is a master regulator of stemness, maintaining self-renewal and pluripotency in embryonic stem cells and various cancers. Its expression is tightly controlled by *SOX2-OT*, a long non-coding RNA that often acts as a positive modulator^123,124^. *SOX2-OT* was uniquely downregulated in MM-LT, suggesting a shift away from stemness and toward myometrial differentiation. Altogether, the SOX gene expression profile in MM LT-cultures highlights pathways related to differentiation, SOX4-mediated contractile activation, tumor suppression and blood vessel formation. The *HOX* transcription factors are master regulators of development, controlling the expression of genes that determine body plan and segment identity during embryogenesis. In mammals, these genes are organized into four clusters (A-D) on different chromosomes^125^. In adult tissues, the cell interaction with the ECM influences *HOX* gene expression and cell phenotype by promoting cell proliferation, adhesion, apoptosis, and migration. Conversely, components of the ECM are also regulated by *HOX* genes, a phenomenon known as dynamic reciprocity^126^. *HOXA* genes are critical regulators of the proper development of the female reproductive tract^127^. *Hoxa11* silencing in the mouse genital tract resulted in alterations of the mRNA and protein expression of types I and III interstitial collagen^128^. Studies on hepatocellular carcinoma have shown that silencing *HOXA11-AS* upregulates *HOXA11*^129^. We found that the antisense *HOXA11-AS* was uniquely upregulated in UL LT-culture, which agrees with the deregulated expression of collagen components frequently observed in UL. We also found *HOXA13* to be highly expressed in UL LT-cultures, confirming previous transcriptomic analysis in *MED12* mutant fibroids^81^. Several genes from clusters B and C, including the lncRNAs *HOTAIR* and *HOXA-AS2*, were uniquely upregulated in MM LT-culture (Table 1). In tissues undergoing regeneration, wound healing, or cyclical renewal (such as skin, endometrium, and bone marrow), *HOX* gene expression can be reactivated to promote proliferation and differentiation^130,131^. The upregulation of *HOX* genes during myometrial repopulation in LT-culture suggests they play a central role in regulating normal MM cell proliferation, migration, and ECM production. Notably, the myometrium undergoes dramatic and reversible structural changes during gestation, parturition, and the postpartum period, and emerging evidence suggests that *HOX* genes may be involved in the dynamic adaptations of the myometrium^126^.

*MYCN* and *MYCL* are members of the MYC proto-oncogene family of transcription factors, which regulate critical cellular processes such as the cell cycle, proliferation, and differentiation. While both genes share overlapping functions, they differ in their expression profiles, transcriptional strength, and cancer associations^132^. In our dataset, *MYCL* was overexpressed in UL LT-culture (Supplementary Dataset S1), whereas *MYCN* (Supplementary Dataset S2) and its downstream effector *MYCT1* (Supplementary Dataset S3) were upregulated in MM LT-culture, suggesting that each tissue engages distinct MYC-driven proliferation programs. This divergence is further supported by the activation of the *SCF-KIT* pathway only in MM LT-culture (Figure 3, Supplementary Dataset S3). SCF (Stem Cell Factor, also known as KITLG) binds to the tyrosine kinase KIT receptor, initiating a signaling cascade that promotes SC survival, proliferation, migration, and differentiation during spermatogenesis^133^. In the mouse myometrium, a classical *in vivo* pulse-chase experiment identified c-Kit⁺ cells near SC. Based on their spatial proximity and temporal emergence, the authors proposed that these c-Kit⁺ cells arise from SCs through asymmetric division, serving as a transient-amplifying population that progresses toward terminal differentiation^134^. Supporting this finding, we detected KIT+ cells only in MM LT-culture, and the Ki67 proliferation marker was observed in a few MM cells. Conversely, most cells in the UL LT-culture strongly stained for Ki-67 (Supplementary Figure S6). This data suggests that after SC activation, MM adopt a more regulated proliferative state compared to the more actively dividing in UL.

Commonly upregulated markers of SC or progenitor cells, such as CD24 and CD73, previously found upregulated in fibroids, were significantly augmented in the UL LT-cultures (Supplementary Dataset S3, Table 1)^15^. Our previous immunohistochemical analysis found strong immunoreaction for CD24 and CD73 in UL LT-culture, whereas faint or no signal was detected in MM LT-culture, respectively (Supplementary Figure S6).

Finally, the activation of p53-regulated pathways in both normal and tumor uterine cells during proliferation and differentiation in LT-cultures may indicate that cells are engaging genome surveillance mechanisms (Supplementary Figure S5, Figure 2). P53 is a transcriptional regulator of genes related to cell cycle control and DNA repair which may help preserve genomic integrity under proliferative conditions. This activation could contribute to the benign nature of uterine leiomyomas and potentially explain their low rate of malignant transformation into leiomyosarcomas.

Comparison of our data with previous transcriptomic dataset from *MED12*-mutated fibroids and matched myometrium^5^ revealed strong concordance in the deregulated genes, with no genes showing opposing expression patterns across the datasets. Notably, 70% of the 20 most dysregulated genes previously identified in *MED12*-mutated fibroids were similarly dysregulated in our analysis. While most findings were consistent, several genes related to stemness showed divergent expression patterns. This discrepancy highlights a key distinction in experimental design: traditional comparisons between fibroids and paired myometrium capture a static view of basal gene expression in tissues with very limited proliferative capacity. Conversely, our study presents a dynamic model, where low oxygen levels reached after LT-culture activated SCs in their physiological niche, inducing their proliferation and differentiation. This resulted in a pronounced upregulation of SC-associated genes, revealing critical regulatory pathways and genes relevant to MM and UL progression that had not been detected before. These findings underscore the exceptional value of LT-cultures of organs as a robust and physiologically relevant platform for studying the development of normal and tumor tissues.

A limitation of our study is its focus on transcriptomic profiling, which lacks direct data on protein expression and function essential for interpreting cellular dynamics. Nevertheless, our conclusions are supported at two levels. First, we validated findings at the protein level using established SC or progenitor cell markers (CD24, CD73, KIT, CD49b) in MM-LT and UL-LT cultures. These protein changes mirrored those observed at the RNA level, reinforcing our transcriptomic results. Second, we consistently confirmed gene expression signatures and pathway differences previously reported between myometrium and leiomyoma. This convergence supports both the robustness and physiological relevance of the long-term organ culture platform. The future implementation of spatial transcriptomics would enable the mapping of gene expression within intact tissue architecture, capturing cell–cell interactions and further refining our understanding of stem cell niches and microenvironmental drivers of leiomyoma growth.

Altogether, our data offer compelling proof of concept for the use of long-term organ cultures as a powerful tool to investigate hypoxia-driven SC activation. This system not only facilitates the study of leiomyoma pathophysiology but also offers a unique opportunity to explore proliferative dynamics in normal myometrium, a process highly relevant to uterine physiology and human reproduction. Moreover, it opens avenues for uncovering novel mechanisms with high translational relevance in regenerative medicine and cancer therapy.

## Supporting information

Supplementary Figure S1. Histological analysis of T0 and long-term culture slices

Supplementary Figure S2. Preservation of the driver MED12 mutation in UL throughout the long-term culture.

Supplementary Figure S3. Significantly overexpressed pathways in uterine leiomyoma (UL) following LONG-TERM CULTURE (LT) compared to baseline (T0).

Supplementary Figure S4. Significantly overexpressed pathways in myometrium (MM) following LONG-TERM CULTURE (LT) compared to baseline (T0).

Supplementary Figure S5. Dot plot illustrating Reactome pathways significantly enriched (q < 0.05) among genes upregulated in long-term (LT) cultured

Supplementary Figure S6. Differential expression of progenitor cell markers in leiomyoma and myometrial tissue slices after long-term culture

Supplementary Dataset S1. Upregulated and downregulated genes in UL T0 vs UL LT-culture

Supplementary Dataset S2. Upregulated and downregulated genes in MM T0 vs MM LT-culture

Supplementary Dataset S3. Upregulated and downregulated genes in UL LT-culture over MM LT-culture

## Supplementary Material

**Supplementary Figure S1.** Histological analysis of T0 and long-term culture slices.

Representative images of leiomyoma and myometrium tissue sections, hematoxylin and eosin stained (H&E) at baseline (day 0) and after 7, 15, 20, 25, and 29 days of culture. Scale bar 100 μm. (B) Tissue distribution of smooth muscle cells (SMC) at long-term culture (LT) of leiomyoma (UL) and myometrium (MM) slices. DES (red) stains SMCs, whereas VIM (green) stains both SMCs and other cell types. DAPI stains the cell nucleus (blue). White arrows indicate VIM+ cells, more abundant in MM LT-culture. Scale bar 100 μm.

**Supplementary Figure S2.** Preservation of the driver *MED12* mutation in UL throughout the long-term culture.

Sequence electropherograms illustrating mutations in exon 2 of *MED12* in the original tumor (T0) and after long-term culture (T20, T25, and T29). Three tumors (L94, L95, and L98) showed the same point mutation, c.131 G>A, in the codon 44 hotspot (highlighted in yellow). The indel mutation of the remaining tumor (L100) consisted of the deletion of seven nucleotides (positions 118–124) and insertion of a cytosine (red circle).

**Supplementary Figure S3.** Significantly overexpressed pathways in uterine leiomyoma (UL) following LONG-TERM CULTURE (LT) compared to baseline (T0).

Hypoxia-related pathways are highlighted in yellow. Pathway enrichment analysis was performed using Gene Ontology (GO) and Kyoto Encyclopedia of Genes and Genomes (KEGG) databases, with significance determined by adjusted p-values (Benjamini– Hochberg correction). The left panel displays the Normalized Enrichment Score (NES) for each pathway, indicating both the direction (upregulation) and magnitude of pathway activity changes. The right panel presents a scatter plot summarizing adjusted p-values, total gene counts per pathway, and the proportion of differentially expressed genes.

**Supplementary Figure S4.** Significantly overexpressed pathways in myometrium (MM) following LONG-TERM CULTURE (LT) compared to baseline (T0).

Hypoxia-related pathways are highlighted in yellow. Pathway enrichment analysis was conducted using Gene Ontology (GO) and Kyoto Encyclopedia of Genes and Genomes (KEGG) databases, with significance determined by adjusted p-values (Benjamini– Hochberg correction). The left panel displays the Normalized Enrichment Score (NES) for each pathway, reflecting both the direction (upregulation) and magnitude of pathway activity changes. The right panel presents a scatter plot summarizing adjusted p-values, total gene counts per pathway, and the proportion of differentially expressed genes.

**Supplementary Figure S5.** Dot plot illustrating Reactome pathways significantly enriched (q < 0.05) among genes upregulated in long-term (LT) cultured slices compared to baseline (T0) in both leiomyoma (UL) and myometrium (MM) tissue slices. Only pathways consistently significant across both tissue types are displayed. Each dot represents an enriched pathway, with size indicating the GeneRatio (proportion of input genes mapped to the pathway) and color reflecting the adjusted p-value (Benjamini–Hochberg correction). Pathways are ranked by statistical significance.

**Supplementary Figure S6.** Differential expression of progenitor cell markers in leiomyoma (UL) and myometrial (MM) tissue slices after long-term culture (LT). **(A)** Immunofluorescence staining of CD49b (ITGA2) (red), KIT (green), and the proliferation marker Ki67 (green) in UL and MM slices at LT-cultures. Nuclei are counterstained with DAPI (blue). Insets show magnified views of marker localization; arrowheads highlight CD49b⁺ cells in the myometrium. **(B)** Immunofluorescence staining of CD24 (green) and CD73 (green) in UL and MM slices at LT-culture. Nuclei are counterstained with DAPI (blue). Insets provide detailed views of marker expression. Scale bar 100 μm.

Adapted from Salas et al., *Biomedicines*, 2022, 10(7), 1542, under CC BY license.

**Supplementary Dataset S1. Upregulated and downregulated genes in UL T0 over UL LT-culture**

**Supplementary Dataset S2. Upregulated and downregulated genes in MM T0 over MM LT-culture**

**Supplementary Dataset S3. Upregulated and downregulated genes in UL LT-culture over MM LT-culture**

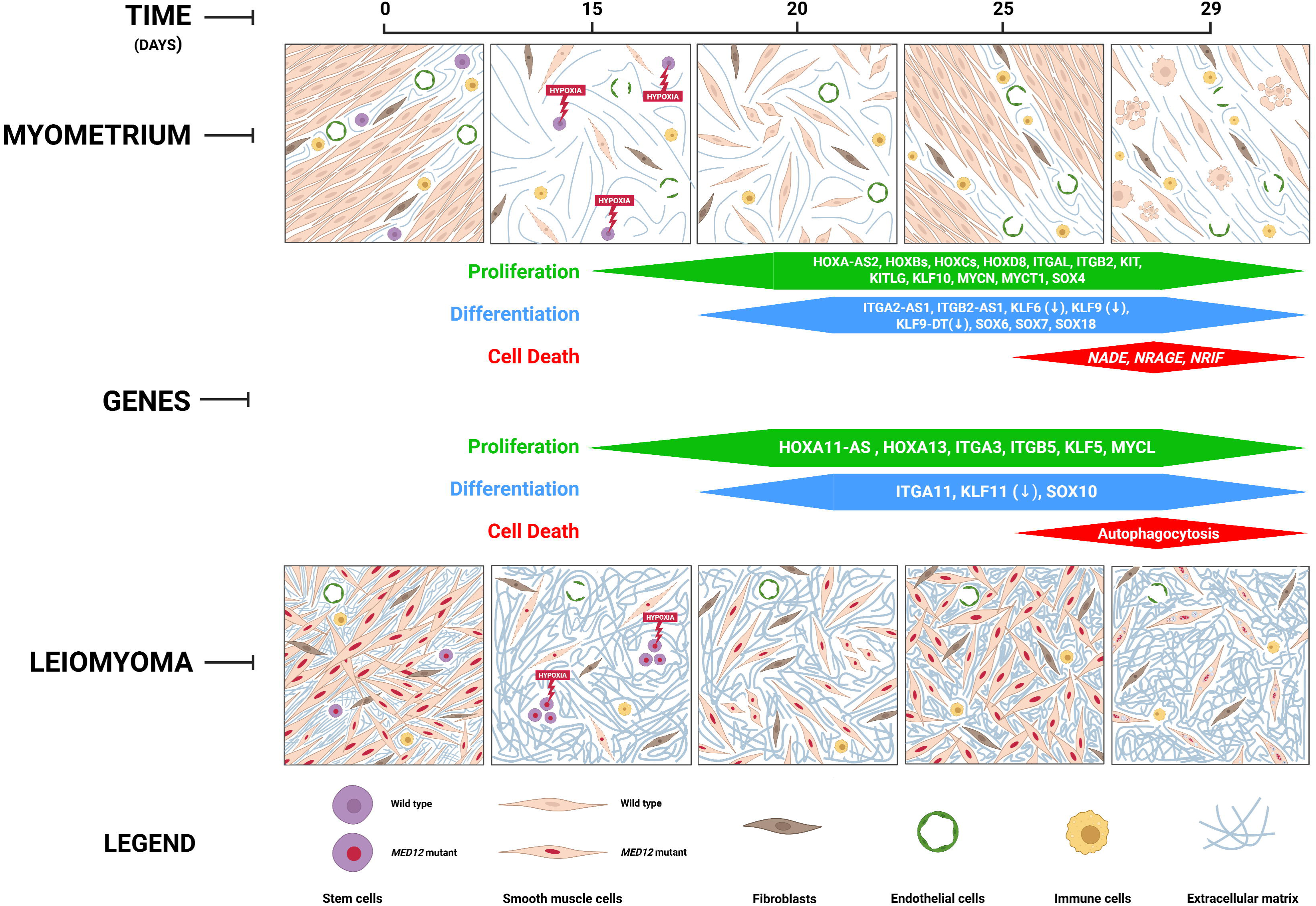

