## Supplementary figures and images for "Long-Term Organ Culture Reveals Differential Stem Cell–Driven Remodeling in Myometrium and *MED12*-Mutant Uterine Leiomyoma"

### Supplementary Figure S1. Histological analysis of T0 and long-term culture slices

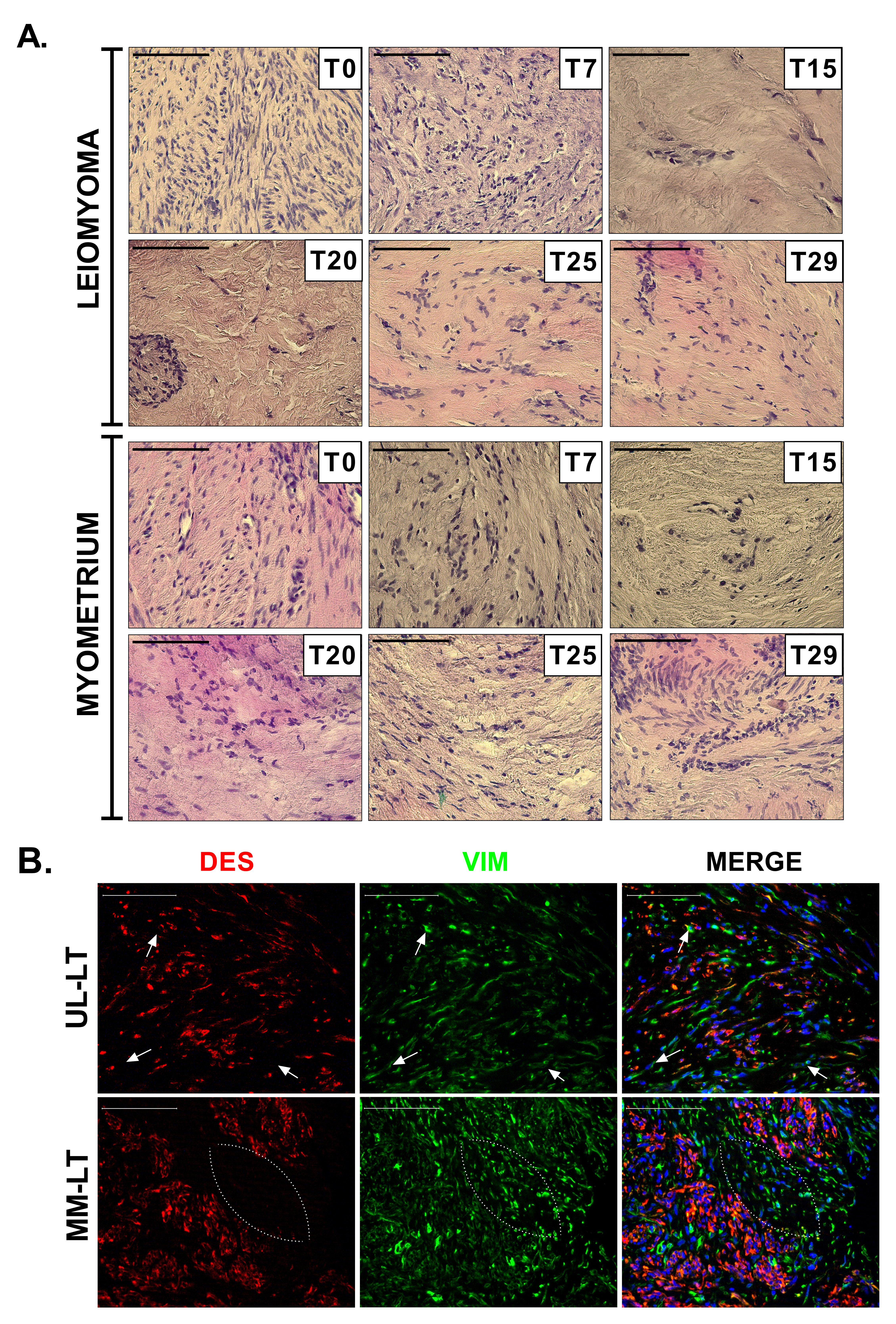

### Supplementary Figure S2. Preservation of the driver MED12 mutation in UL throughout the long-term culture.

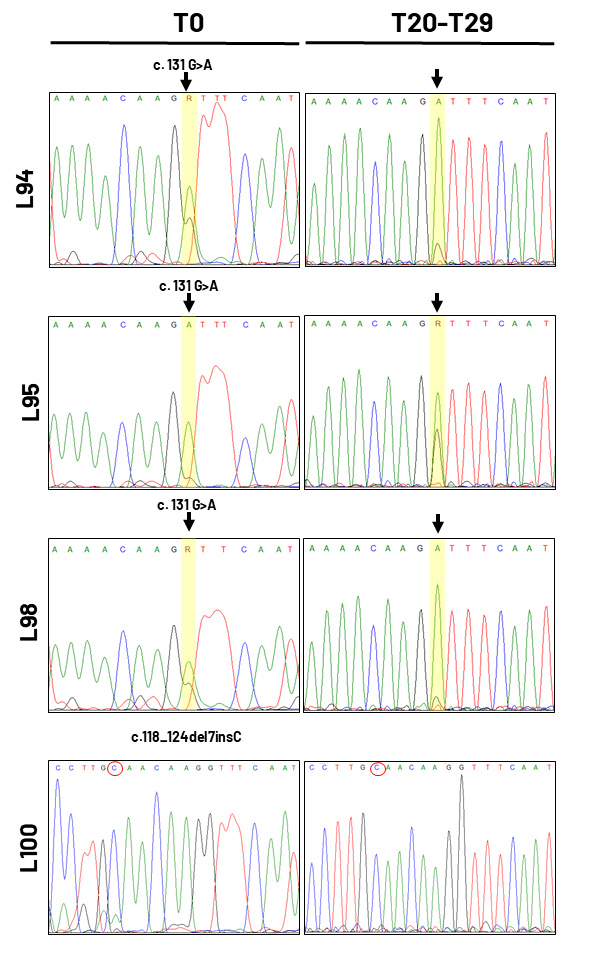

### Supplementary Figure S3. Significantly overexpressed pathways in uterine leiomyoma (UL) following LONG-TERM CULTURE (LT) compared to baseline (T0).

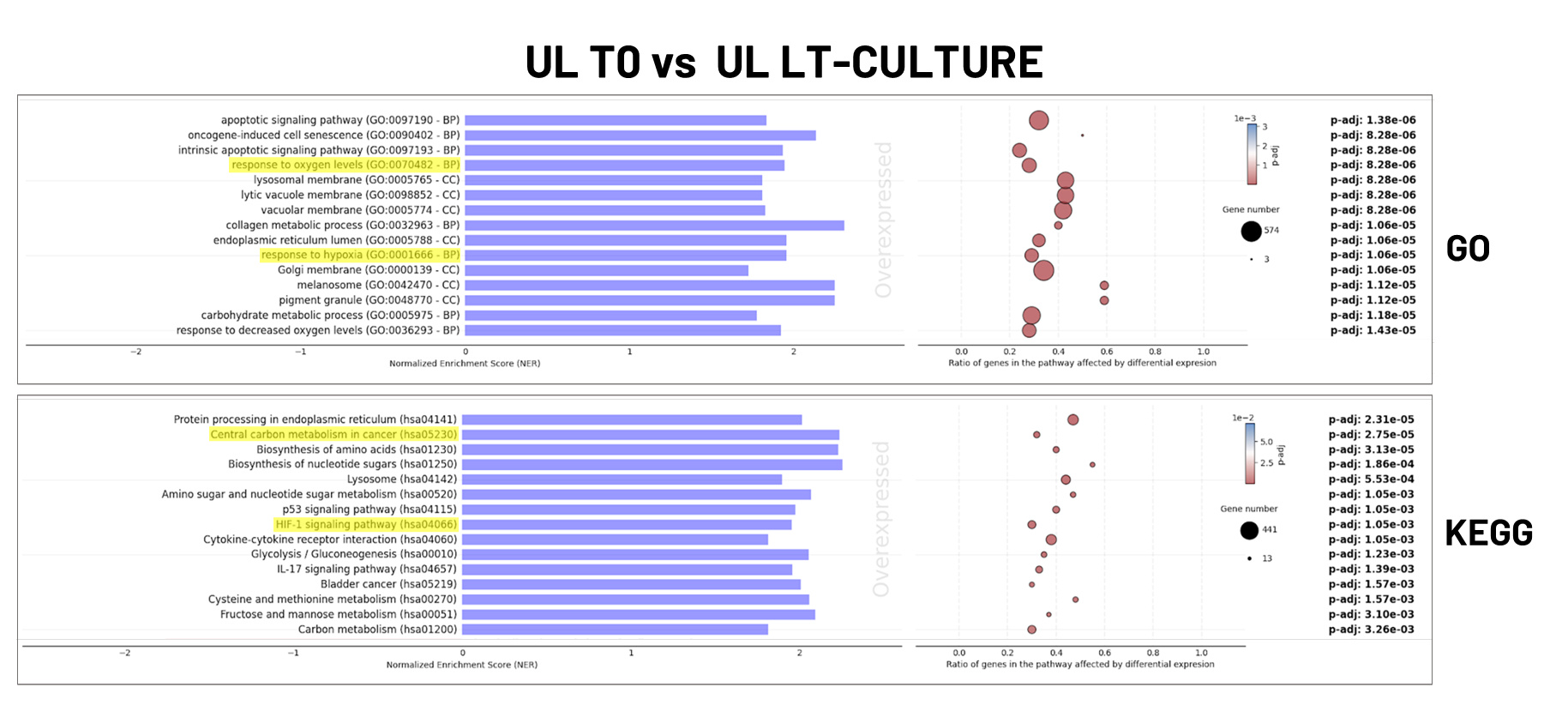

### Supplementary Figure S4. Significantly overexpressed pathways in myometrium (MM) following LONG-TERM CULTURE (LT) compared to baseline (T0).

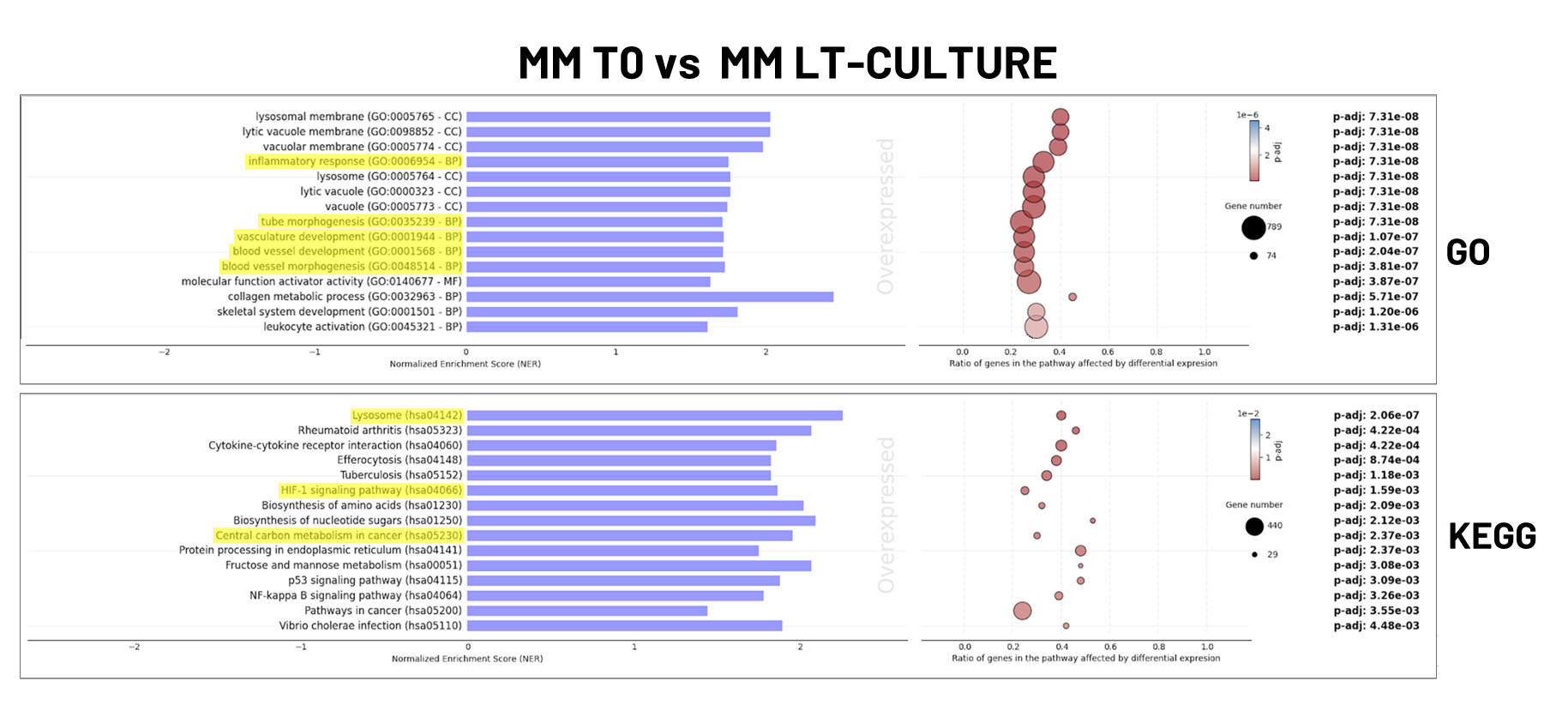

### Supplementary Figure S5. Dot plot illustrating Reactome pathways significantly enriched (q < 0.05) among genes upregulated in long-term (LT) cultured

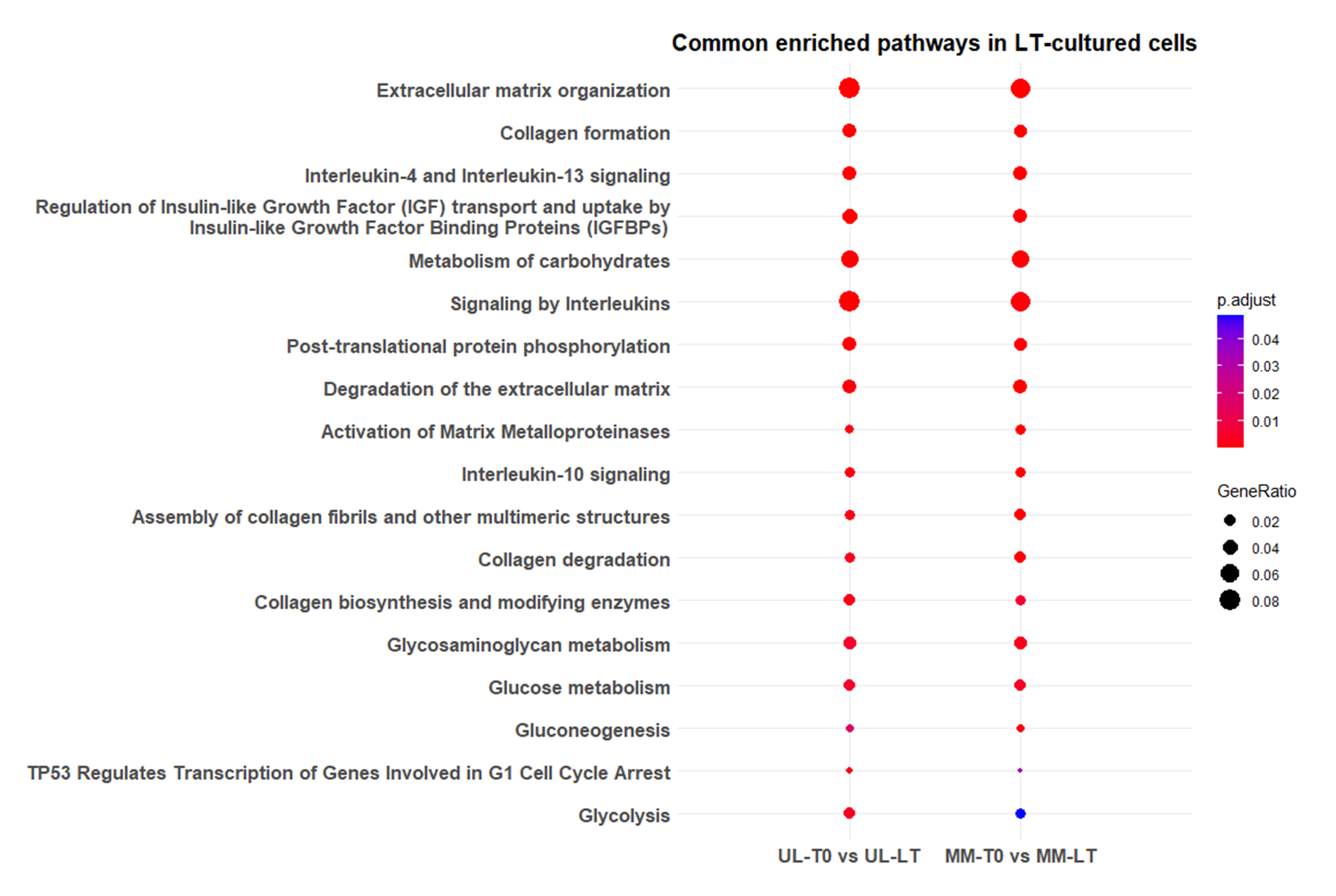

### Supplementary Figure S6. Differential expression of progenitor cell markers in leiomyoma and myometrial tissue slices after long-term culture

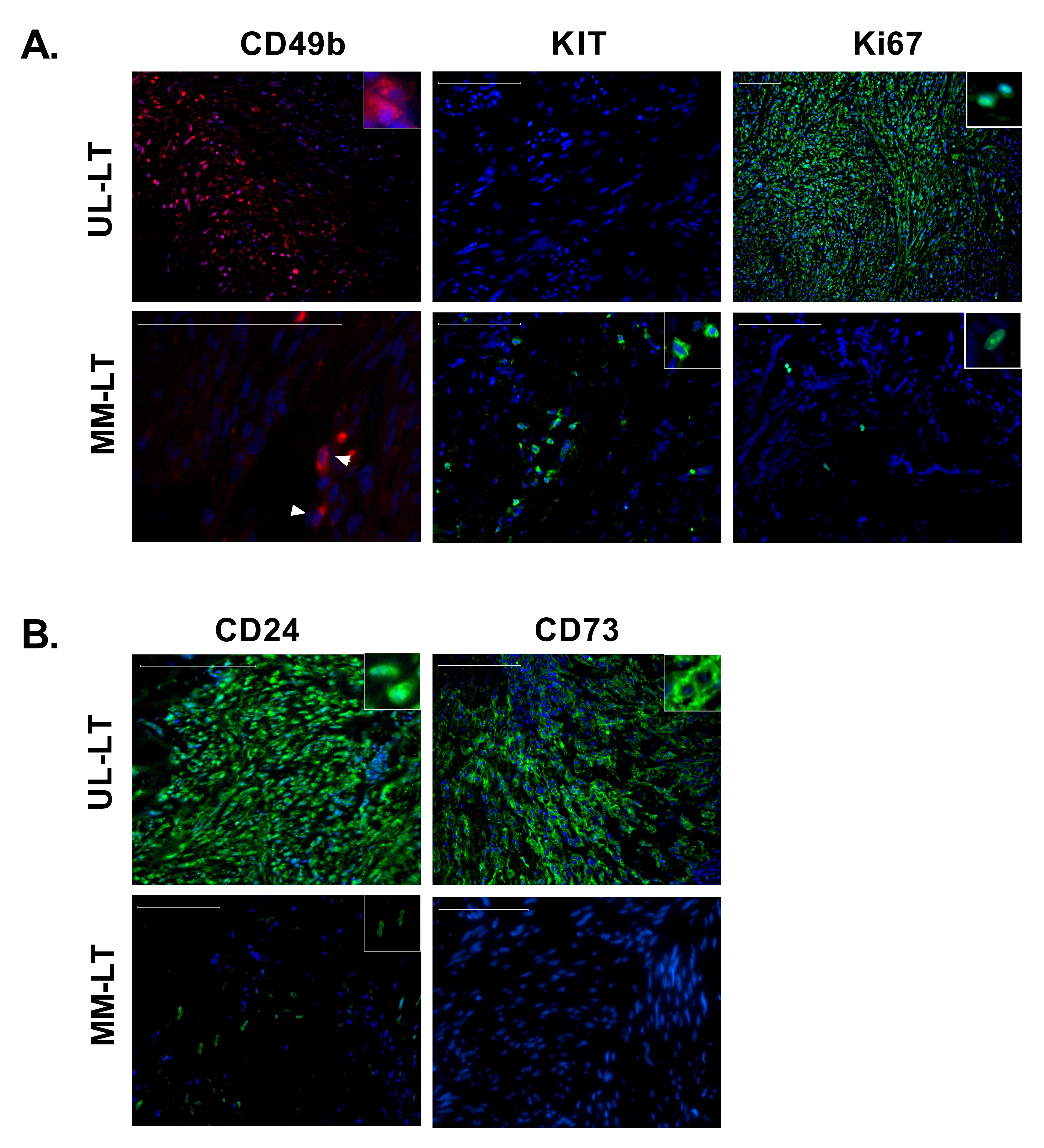
